# Unveiling the role of yeast cytochrome *c* isoforms in the assembly of mitochondrial supercomplexes and the control of respiratory chain rate

**DOI:** 10.1101/2024.07.13.603375

**Authors:** Alejandra Guerra-Castellano, Manuel Aneas, Joaquín Tamargo-Azpilicueta, Inmaculada Márquez, José Luis Olloqui-Sariego, Juan José Calvente, Miguel A. De la Rosa, Irene Díaz-Moreno

## Abstract

Mitochondria play crucial roles as both the powerhouse and signaling center of cells, balancing cell survival and death to maintain homeostasis. Disruption of this balance can lead to various diseases. Therefore, exploring the components involved in mitochondrial metabolism presents a significant challenge. In this context, respiratory supercomplexes are evolutionarily conserved, stable associations between membrane complexes and molecules, including proteins and lipids, within the inner mitochondrial membrane. These supercomplexes dynamically respond to metabolic demands, enhancing the electron transfer rate and reducing the production of reactive oxygen species. Recent research has identified cytochrome *c*, a mobile electron carrier between complexes III and IV, as a potential key player in the formation of these supercomplexes. This study focuses on elucidating the role of cytochrome *c* in modulating the assembly of supercomplexes, using the yeast *Saccharomyces cerevisiae* as a model system for mitochondrial metabolism. Our findings indicate that the viability of Saccharomyces cerevisiae relies on the presence of cytochrome *c*, with both isoforms playing a role in the assembly of respiratory supercomplexes. Notably, isoform-2 of cytochrome *c* enhances electron transfer efficiency, resulting in reduced ROS production.

## 1. Introduction

Mitochondria are the main source of reactive oxygen species (ROS) in the cell, where the complex I (CI) and complex III (CIII) are the principal entities that generate them (Pizzino *et al*., 2017). Cells rely on several mechanisms to reduce the impact of ROS, through the production of antioxidant molecules or by rearranging the electron transport chain (ETC) elements into high-ordered structures (Poljsak, 2011).

The ETC supramolecular organization has been discussed in detail, and three possible models have been proposed. The *solid* model claims a steady assembly of the respiratory complexes into a high-order molecular structure named oxysome, whereas the *fluid* model considers the free and independent diffusion of the mitochondrial complexes within the inner mitochondrial membrane (IMM) (Chance & Williams, 1955; Hackenbrock *et al*., 1986; Schägger & Pfeiffer, 2000; Lenaz & Genova, 2007, 2012). Finally, the *plasticity* model suggests a variety of respiratory complex associations depending on metabolic needs (Acín-Pérez *et al*., 2008; Acín-Perez & Enriquez, 2014; Letts *et al*., 2016).

The respiratory supercomplexes are higher associations, which can contain complex I, II, III, and IV (CI, CII, CIII, and CIV, respectively) and, rarely, complex V (CV) (Vartak *et al*., 2013). These assemblies reduce the distance between the components of the ETC, therefore they enhance the efficiency of electron transport, and reduce ROS production (Maranzana *et al*., 2013; Genova & Lenaz, 2014; Brzezinski *et al*., 2021). Supercomplexes have been isolated from several organisms (mammals, plants, yeasts, and bacteria), and their structures have been resolved by cryo-electron microscopy (Stroh *et al*., 2004; Gu *et al*., 2016; Letts *et al*., 2016; Sousa *et al*., 2016; Wu *et al*., 2016; Guo *et al*., 2017; Gong *et al*., 2018; Wiseman *et al*., 2018; Berndtsson *et al*., 2020; Maldonado *et al*., 2021; Steimle *et al*., 2021; Moe *et al*., 2022). Remarkably, the stability of the individual complexes in the membrane is not affected by the supercomplexes assembly, since their levels are not altered in the absence of supercomplexes (Cruciat *et al*., 2000).

The supercomplexes formation implicates specific interactions between the respiratory complexes that are mediated and/or modulated by several proteins and lipids (Zhang *et al*., 2005; Dienhart & Stuart, 2008; Claypool, 2009; Strogolova *et al*., 2012; Vukotic *et al*., 2012; Ikeda *et al*., 2013; Vartak *et al*., 2013; Friedrich *et al*., 2016; Singhal *et al*., 2017; Shimada *et al*., 2018; Zong *et al*., 2018; Díaz-Quintana A *et al*., 2020; Timón-Gómez *et al*., 2020). Recently, the mobile carriers quinone (Q) and cytochrome *c* (C*c*) could mediate or modulate the interaction of supercomplexes because it has been reported a comigration of them in native gels with different supercomplex associations (Acín-Pérez *et al*., 2008), and the absence of C*c* bring lack of CIV assembly (Vempati *et al*., 2009). These two findings highlight that C*c* might play a role as a supercomplex assembly factor of mitochondrial respiratory complexes’ assembly and its association.

To delve deeper into this aspect, the yeast *Saccharomyces cerevisiae* (*S. cerevisiae*) has been used in this study because has been extensively used as a model organism thanks to its genetic amenability. Its genome was one of the first eukaryotic genomes sequenced (Goffeau *et al*., 1996). It shares many conserved cellular pathways with higher eukaryotes, including humans. Therefore, discoveries in yeasts may be extrapolated to comprehend similar processes in other organisms and are very useful in studying some human diseases (Botstein *et al*., 1997; Nielsen, 2019). In addition, this yeast strain is easy to grow, showing an energy metabolism depends on current oxygen concentration and carbon source. Concerning ETC, *S. cerevisiae* does not harbor complex I (CI). Instead of that, it has three membrane peripheral NADH dehydrogenases. Two of them are located at the outer surface (Nde1p and Nde2p) and the other one at the inner surface (Ndi1p) of the inner mitochondrial membrane (IMM) (Luttik *et al*., 1998; Matus-Ortega *et al*., 2015) (Figure 1). These dehydrogenases do not pump protons through the IMM, i.e., they do not contribute to the electrochemical gradient (Bakker *et al*., 2001). Due to the absence of CI, *S. cerevisiae* shows two types of supercomplexes association, composed of a CIII dimer which is enclosed by either one or two CIV (III_2_IV_1_ and III_2_IV_2_, respectively) (Schägger and Pfeiffer, 2000; Hartley *et al*., 2019; Rathore *et al*., 2019; Berndtsson *et al*., 2020; Hartley *et al*., 2020; Moe *et al*., 2021).

**Figure 1.**
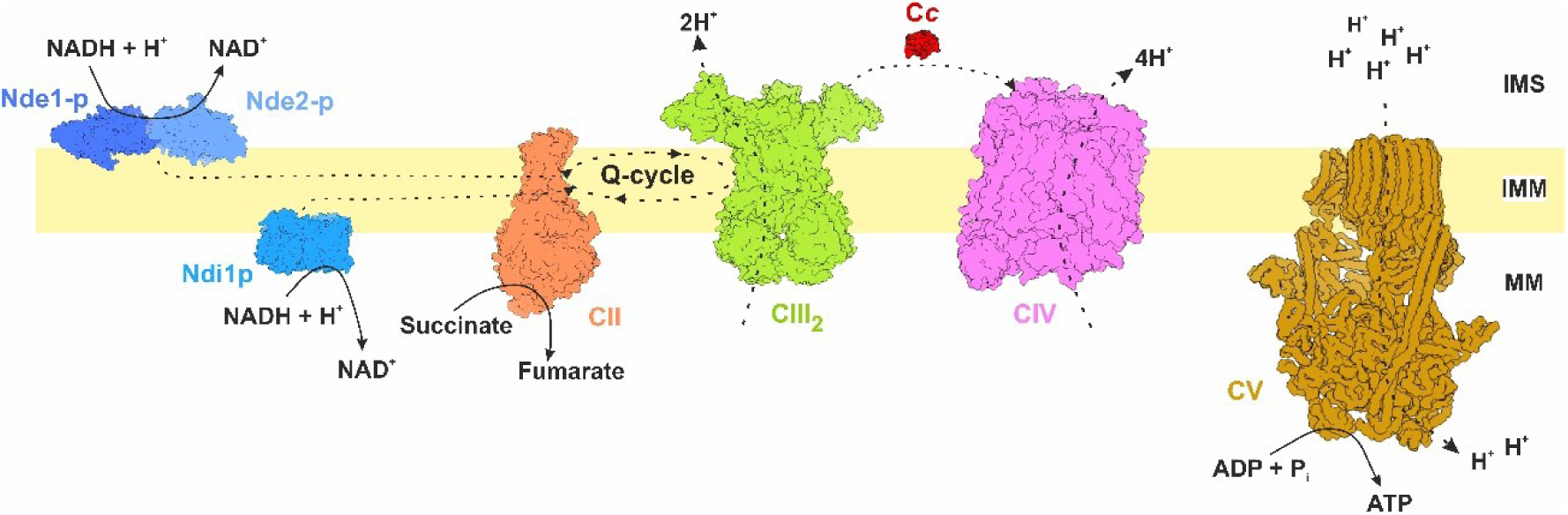
Overview of oxidative phosphorylation and electron transport chain in *Saccharomyces cerevisiae*. The OXPHOS system in *S. cerevisiae* is similar to the mammalian system, with the exception that complex I is replaced by a non-proton-translocating NADH dehydrogenase (Ndi1p) situated on the inner side of the IMM. Moreover, *S. cerevisiae* has two NADH dehydrogenases (Nde1p, Nde2p) on the outer side of the IMM, which transfers electrons to ubiquinone. CII: complex II, CIII: complex III, CIV: complex IV, CV: complex V, C*c*: iso-1 cytochrome *c*, IMS: Inter mitochondrial space, IMM: inner mitochondrial membrane, MM: mitochondrial matrix, Q: quinones. All structures represented are from *S. cerevisiae*, except CII which is from human. Nde1-p: Alphafold P40215; Nde2-p: Alphafold Q07500; Ndpi1p: PDB 5YJY (Yamashita *et al*. 2018); CII: PDB 8GS8 (Du *et al*. 2023); CIII: PDB 1KYO (Lange *et al*. 2002); CIII: PDB 6YMY (Berndtsson *et al*., 2020); CV: PDB 5VOY (Zhao *et al*. 2017). Figure has been generated with software ChimeraX (Meng *et al*. 2023).

*S. cerevisiae* has two isoforms of C*c* namely iso-1 and iso-2. These two isoforms are encoded by two different genes: *cyc1* and *cyc7*, (iso-1 and iso-2 C*c*, respectively). The iso-1 C*c* is the homolog of the human C*c*, which replaces iso-1 C*c*in yeasts (Ying *et al*., 2009). The C*c* isoforms are found in a 95:5 ratio (iso-1:iso-2) under normoxic conditions, whereas in hypoxic conditions the iso-2 C*c* prevails (Burke *et al*., 1997; Kwast *et al*., 1999). The isoforms share 84% of sequence identity, and the iso-2 C*c* has a four-residue extension in the N-terminus (Figure S1 in Appendix A) (Downie *et al*., 1977; Montgomery *et al*., 1980). To date, the biological meaning of hosting two C*c* isoforms in yeasts is unknown, considering that *S. cerevisiae* primarily undergoes fermentation. This study aims to investigate the function of these isoforms in mitochondrial metabolism and their potential involvement in assembling respiratory supercomplexes.

## 2. Material and methods

### 2.1. Cloning, expression, and purification of recombinant cytochrome *c* isoforms

To obtain both C*c* isoforms from yeast *S. cerevisiae*, *Escherichia coli* BL-21 (DE3) cells were transformed with the plasmids pBTR1-cyc1 and pBTR1-cyc7 —which contains the coding region for the iso-1 C*c* and iso-2 C*c*, respectively— and the *cyc3* gene that encodes the yeast C*c* heme lyase, which is necessary for the proper maturation of the heme group (Olteanu *et al*., 2003). Plasmids pBTR1-cyc1 and pBTR1-cyc7 were obtained by replacing the *cycs* gen (which encoded for human C*c*) with the yeast *cyc1* or *cyc7* genes, respectively. Yeast genes have been extracted from *S. cerevisiae* nuclear DNA by performing a PCR with the primers described in Table S1 (Appendix A).

After transformation, cells were incubated for 24 h at 30 °C and 150 rpm in Luria-Bertani (LB) medium (10 g/L tryptone, 5 g/L yeast extract, 10 g/L NaCl) supplemented with ampicillin (final concentration of 0.1 mg/mL). Cells were collected by centrifugation and then resuspended in lysis buffer (10 mM Tricine-NaOH, pH 8.5, supplemented with 1 mM phenylmethylsulfonyl fluoride (PMSF), 0.02 mg/mL DNase, and 0.2 mg/mL complete protease inhibitors). The cytoplasmic fraction was obtained by sonication followed by centrifugation at 20,000 *g* for 15 min. The supernatant was collected and loaded onto a Nuvia-S column (Bio-Rad, Hercules, CA, USA) using a fast protein liquid chromatography (FPLC) system (Bio-Rad). Proteins were eluted from the column using a NaCl gradient (0-1 M), and fractions containing the isoforms were identified by UV-Vis spectrometry (A_550_). The purity of the proteins was analyzed using the A_280_/A_550_ ratio (*ca* 1.0) of the resulting reduced C*c* preparations, as previously described (Guerra-Castellano *et al*., 2015). The concentration of the recombinant proteins was determined by visible spectrophotometry using the molar extinction coefficient for reduced C*c* species (ε = 28 mM^-1^·cm^-1^).

### 2.2. Mass spectrometry

A mass spectrometry analysis was performed to confirm C*c* isoforms. A linear positive mode MALDI-TOF (Matrix-Assisted Laser Desorption/Ionization Time-of-Flight) Ultraflextreme (Bruker) spectrometer was utilized. The results were averaged from 3,000 laser shots in each spectrum. A 1:10 mixture of α-cyano-hydroxycinnamic acid and 2,5-dihydroxybenzoic acid solutions (20 mg/mL) was used as the MALDI matrix. This matrix was mixed with each of C*c* isoform samples (iso-1 and iso-2) in a 1:2 volume ratio, and 0.5 μL was deposited onto the sample holder on an α-cyano-hydroxycinnamic acid film. The deposited droplet was left to dry at room temperature. The spectra were recorded at the Biomolecular Mass Spectrometry Service of Pablo de Olavide University.

### 2.3. Gel electrophoresis under denaturing conditions

An electrophoresis in polyacrylamide gel (15% acrylamide/bisacrylamide) was carried out under denaturing conditions (SDS-PAGE) to corroborate the purity of preparations. 10 μg of each C*c* isoform and 5 μL of Tris-Glycine 4-20% BlueStar PLUS Prestained Protein Marker (Nippon Genetics) were loaded into the gel. It was stained for protein detection with 0.25% Coomassie Brilliant Blue R-250 in a solution of 45% methanol and 10% acetic acid.

### 2.4. Tryptic digestion

The bands of each C*c* isoform of the SDS-PAGE gel were subject to a tryptic digestion assay. Acrylamide gel bands were detained with ammonium bicarbonate and acetonitrile. Dithiothreitol (DTT) and iodoacetamide were respectively used to break disulphide bonds and carbamidomethylate cysteine residues. Samples were incubated overnight at 37 °C with bovine trypsin at ratio 1:10 (enzyme:substrate). After acetonitrile extraction and acidification, samples were desalted and concentrated with C18-filled tips.

Tryptic digestion peptides were separated using a 50cm C18 EASY-Spray column coupled with a C18 PepMap100 2 cm pre-column. A non-linear gradient was used between phase A (0.1% formic acid) and phase B (80% acetonitrile, 20% water, 0.1% formic acid), during 120 min in reverse phase mode. A Thermo ScientificTM Q Exactive™ Plus Orbitrap™ mass spectrometer was used to acquire the top 10 MS/MS spectra in DDA mode. Spray Voltage was fixed at 2.9 kV, and capillary temperature at 300 °C. Maximum IT was set at 30 ms.

Liquid chromatography–mass spectrometry (LC-MS) data were analyzed using SEQUEST® HT search engine, in Thermo Scientific™ Proteome Discoverer™ 2.2 software. The following aminoacid modifications have been considered in the analysis: static carbamidomethylation (C), dynamic oxidation (M) and dynamic acetylation (N-terminal). Data were searched against specific protein sequences for each mutant and results were filtered using a 0.01% protein FDR threshold.

The spectra were recorded at the Biomolecular Mass Spectrometry Service of the Pablo de Olavide University using a MALDI-TOF Ultraflextreme spectrometer (Brucker).

### 2.5. Circular dichroism spectroscopy

Circular dichroism (CD) spectra were recorded using a Jasco® J-815 spectropolarimeter equipped with a Peltier temperature-control system.

Secondary structure analysis of C*c* isoforms was carried out by recording far UV (185 – 250 nm) CD spectra at 25 °C in a 1 mm quartz cuvette. Samples of each protein were analyzed separately, with a concentration of 3 μM protein in 10 mM sodium phosphate (pH 6.5), supplemented with 10 μM potassium ferrocyanide. For each sample, 20 scans were averaged.

The coordination of the heme iron atom of the S_δ_ atom of methionine was analyzed by recording Vis (300 – 600 nm) CD spectra at 25 °C in a 10 mm quartz cuvette (Blauer *et al*., 1993; Guerra-Castellano *et al*., 2016). As before, samples of each protein were analyzed separately, with a concentration of 30 μM protein in 10 mM sodium phosphate (pH 6.5), supplemented with 10 μM potassium ferrocyanide.

### 2.6. Electrochemical measurements

The linear scan voltammetric experiments were performed using an AUTOLAB PGSTAT 30 system (Eco Chemie B.V.), in a three-electrode undivided glass cell equipped with a gas inlet and a temperature control system. The working electrodes were polycrystalline gold with a geometric area of 0.0314 cm^2^.

Before measurements, the gold surface was cleaned by successive polishing with 0.3 and 0.05 μm alumina, rinsed with Millipore water, and sonicated in absolute ethanol to remove residual alumina. Then, the electrode surface was dried and chemically cleaned using a "piranha" solution (7:3 concentrated H_2_SO_4_ (95%) and 30% (v/v) H_2_O_2_). Gold electrodes were modified with thiol self-assembled monolayers (SAMs) by immersing the electrode in ethanol solutions containing 1 mM 8-mercaptooctanoic acid (MOA) and 2.5 mM 8-mercapto-1-octanol (MOOL) (Sigma-Aldrich Co.) for 30 min at room temperature. Protein immobilization was carried out by depositing a 15 µL drop of a solution containing 20 µM of protein and 10 mM sodium phosphate buffer (pH 7) onto the modified electrode for 45 min at 4 °C. After protein incubation, the electrodes were rinsed with water and washed with the working buffer. The counter and reference electrodes were a platinum bar and an Ag/AgCl/NaCl-saturated electrode, respectively. The reference electrode was connected to the cell solution via a salt bridge and kept at room temperature (298 ± 2 K) in a non-isothermal configuration. The obtained potential values were corrected to the normal hydrogen electrode (NHE) potential scale by adding + 192 mV to the experimental values. The working solutions contained a 20 mM sodium phosphate buffer as the supporting electrolyte at pH 7.0 ± 0.1. All measurements were performed under an argon atmosphere (Olloqui-Sariego *et al*., 2022).

Thermodynamics and kinetics of the electron exchange between protein and electrode were obtained from the analysis of the voltammetric peak potentials as a function of temperature and potential scan rate, respectively. To this end, temperature variable cyclic voltammetry was performed at equispaced temperatures, whose upper limit was dictated by protein thermal desorption. A positive feedback for ohmic drop compensation was applied when the potential scan rate of the voltammograms was higher than 1 V s 1.

### 2.7. Yeast strains and growth conditions

*S. cerevisiae* strains used in this study are the wild-type (WT; BY4741; *MAT*a; *his3*Δ*1*; *leu2*Δ*0*; *met15*Δ*0*; *ura3*Δ*0*), iso-1 C*c* deficient (Δ*cyc1*: BY4741; *MAT*a; *his3*Δ*1*; *leu2*Δ*0*; *met15*Δ*0*; *ura3*Δ*0*; *YJR048w::kanMX4*), iso-2 C*c* deficient (Δ*cyc7*: BY4741; *MAT*a; *his3*Δ*1*; *leu2*Δ*0*; *met15*Δ*0*; *ura3*Δ*0*; *YEL039c::kanMX4*), iso-1 C*c* complementary (*cyc1*^+^: BY4741; *MAT*a; *his3*Δ*1*; *leu2*Δ*0*; *met15*Δ*0*; *ura3*Δ*0*; *YJR048w::kanMX4*; pEC189-*cyc1*) and iso-2 C*c* complementary (*cyc7*^+^: BY4741; *MAT*a; *his3*Δ*1*; *leu2*Δ*0*; *met15*Δ*0*; *ura3*Δ*0*; *YEL039c::kanMX4*; pEC189-*cyc7*). The mutant strains come from the EUROSCARF repository (Δ*cyc1*: Y06846 and Δ*cyc7*: Y00280, respectively). On the other hand, to obtain the complementary yeast strains, the deficient strains were transformed with the plasmids pCM189-*cyc1* and pCM189-*cyc7*, which contain the genes *cyc1* and *cyc7*, respectively (Saad *et al*., 2014). The primers utilized are listed in Table S2 (Appendix A).

The yeast strains were cultured in YPD (10 g/L Yeast extract, 20 g/L Peptone, and 20 g/L Dextrose) or YPG (10 g/L Yeast extract, 20 g/L Peptone, and 3% (v/v) Glycerol) media to study the mitochondrial metabolism dependent on the presence of supercomplexes. The culture conditions were 30 °C, 200 rpm, for 24 h or 48 h (YPD or YPG, respectively). The YPD medium is composed of 10 g/L yeast extract, 20 g/L peptone, and 2 % (w/v) glucose, and the YPG medium contains 10 g/L yeast extract, 20 g/L peptone, and 3 % (v/v) glycerol. For growth patterns assay, these media were supplemented with agar at a final concentration of 20 g/L.

### 2.8. Mitochondria isolation

Mitochondria were isolated from the strains (WT, Δ*cyc1*, Δ*cyc7*, *cyc1*^+^, and *cyc7*^+^), which were culture in YPD and YPG media as described before (Moreno-Beltrán *et al*., 2017). Cells were resuspended in a pretreatment buffer (100 mM Tris-SO_4_, pH 9.4, 10 mM DTT) and were incubated at 34 °C and 200 rpm for 20 min. After this, cells were collected by centrifugation at 1,000 g for 5 min and were resuspended in a digestion buffer (1.2 M sorbitol, 20 mM KP_i_, pH 7.4) to incubate them at 34 °C and 200 rpm for 60 min. Once incubation was completed, centrifugation was performed at 1,000 g for 5 min at 4 °C. The collected cells were resuspended in a lysis buffer (0.6 sorbitol, 20 mM K-MES, pH 6.0) and 100 mM PMSF. Then they were lysed using a Dounce homogenizer. After lysis, centrifugation was carried out at 2,000 g for 5 min at 4 °C, and the pellet was discarded. The supernatants were centrifugated at 12,000 g for 15 min at 4 °C to precipitate mitochondria. The collected mitochondria were stored at – 80 °C in a buffer composed of 0.6 M sorbitol and 20 mM 2-(N-morpholino)ethanesulfonic acid, 4-morpholinoethanesulfonic acid (MES)/KOH, pH 6.0 (Padilla-López *et al*., 2009).

### 2.9. Gel electrophoresis under native conditions

An electrophoresis gel was carried out in native conditions (BN-PAGE) as described previously (Moreno-Beltrán *et al*., 2017). The isolated mitochondria were permeabilized and loaded into the gel. For each strain, 400 μg of mitochondria were permeabilized in 40 μL of solubilization buffer (30 mM HEPES buffer (pH 7.4), with 150 mM KOH-acetate, 10 % glycerol, and 1 mM PMSF) supplemented with digitonin at a digitonin:protein ratio of 4:1 (w:w). Samples were incubated on ice for 30 min and then were centrifugated for 30 s to eliminate the membrane fraction. The supernatants were loaded into a protein gel NativePAGE Novex 3-12 % Bis-Tris (1.0 mm, 15-well; Thermo Fisher Scientific, catalog number BN2012BX10), following the manufacturer’s instructions.

### 2.10. Cytochrome *c* oxidase activity

Two different assays have been performed to measure C*c*O activity: with isolated proteins (*in vitro*) and in mitochondria context (*in mitochondria*). Cytochrome *c* Oxidase Activity Assay kit (BioChain®, catalog number KC310100) has been used for both measurements following the manufacturer’s instructions in a Jasco® V-650 spectrophotometer.

*In vitro* assay has been carried out with bovine CIV and both C*c* isoforms, while *in mitochondria* assay has been performed with isolated mitochondria from all described yeast strains which have been cultured in YPD and YPG media. The detergent n-dodecyl-β-D-maltoside (βDDM) was used to permeabilize the OMM and allow the entry of exogenous C*c* (iso-1 and iso-2) into the mitochondria, following previously described protocols (Musatov *et al*., 2000; Moreno-Beltrán *et al*., 2017). The absence of oxidation signals from endogenous C*c* compels the need to introduce exogenous C*c* to assess C*c*O activity within mitochondria. The C*c*O activity values have been calculated from slopes of absorbance at 550 nm from at least three independent measurements.

### 2.11. Reactive oxygen species measurements

ROS production was measured in isolated mitochondria as previously described (Starkov, 2010; Guerra-Castellano *et al*., 2018). Mitochondria were isolated from yeast strains described in Section 2.7, which have been cultured in YPD and YPG media. Mitochondria (final concentration of 0.06 mg/mL) were incubated in a ROS buffer (125 mM KCl, 4 mM KH_2_PO_4_, 14 mM NaCl, 20 mM HEPES-NaOH, pH 7.2, 1 mM MgCl_2_, 0.2 % of fatty acids free bovine serum albumin, and 0.020 mM EGTA), supplemented with 0.08 mM βDDM to permeabilize OMM for 5 min at 4 °C. Then, fluorescence emitted by the fluorogenic substrate Amplex^TM^ Red (Thermo Fisher Scientific) was recorded on a Cary Eclipse spectrofluorometer (Varian) using an excitation wavelength of 555 nm and an emission wavelength of 581 nm for 10 min. The assay consists of two steps, in the first one 10 μM Amplex^TM^ Red was added, and fluorescence was recorded for 2 min. Then, 10 mM succinate and 4 U/mL horseradish peroxidase were added, and fluorescence was recorded for 4 min. Succinate was used as NAD^+^ dependent-substrate and antimycin A as CIII inhibitor.

### 2.12. Molecular Dynamics simulations

Yeast iso-1 (tri-methylated at Lys77) and iso-2 C*c* (tri-methylated at Lys81) structures were fetched from PDB accessions 3CX5 (Solmaz *et al*., 2008) and 3CXH (Solmaz *et al*., 2008), respectively. The tri-methylated residues were replaced by unmodified Lys using a similar rotamer configuration. Setup was performed using the leap module of AmberTools23 package (Case *et al*., 2023). The force field parameters of the ferrous heme group, His, Cys and Met ligands, were already available at our group (Gomila *et al*., 2022), which were adapted from the AMBER parameters database (http://research.bmh.manchester.ac.uk/bryce/amber) using restrained electrostatic potential (RESP) charges from Autenrieth *et al*. (Autenrieth *et al*., 2012). Force field parameters of tri-methylated lysine were derived from Papamokos *et al*. (Papamokos *et al*., 2012). The system was defined with the ff14SB force field (Maier *et al*., 2015) and solvated in an orthorhombic box of optimal 3-charge, 4-point (OPC) rigid water model molecules. Minimum distance of box edges to the protein surface was set to 10 Å. The net charge of the system was neutralized with chlorine and sodium counter-ions. A 5 ns relaxation step preceded 1 μs molecular dynamics (MD) simulations. Both steps were performed with OpenMM 8.0 software (Eastman *et al*., 2024) in a CUDA platform. Temperature was kept constant at 298 K and controlled by a Langevin thermostat, with a friction coefficient of 1 ps^-1^ and a step size of 0.002 ps. Particle mesh Ewald was set to compute long-range interactions with a direct space cutoff of 10 Å and an error tolerance of 5 · 10^-4^ kJ · mol^-1^, and constraints to the length of bonds that involve a hydrogen atom were applied. For each construct, three replicas were simulated and analyzed. The trajectory analysis was performed with the cpptraj module of AmberTools23 (Roe *et al*., 2013). Graphic representation and further analysis were performed with OriginPro Version 2024.

## 3. Results and Discussion

### 3.1. Recombinant cytochrome *c* isoforms expression enables the study of their biophysical, structural, and electrochemical properties

Isoforms 1 and 2 of *S. cerevisiae* C*c* have been successfully expressed and purified. An SDS-PAGE was run to confirm the homogeneous production of the recombinant proteins (Figure S2A in Appendix A). The bands in the gel were submitted to tryptic digestion assay to verify each isoform. The obtained peptides were analyzed and compared to the sequences of each isoform (Figure S2B in Appendix A, and Appendix B), albeit no peptides corresponding to the N-end extension of iso-2 were detected. Moreover, the isoforms’ molecular masses were obtained and compared with the theoretical them (12671.97 Da and 13022.31 Da, iso-1 and iso-2 C*c*, respectively) (Figure S2C in Appendix A). The largest size of iso-2 C*c* calculated by mass-spectrometry perfectly agrees with the less mobility of the protein in SDS-PAGE

Once we have confirmed the C*c* isoforms production, their folding and heme group coordination were analyzed by CD spectroscopy. Far-UV CD spectra give information about the secondary structure elements due to it is sensitive to the peptide bonds. The main secondary structure in both isoforms was α-helix, as expected for C*c* proteins (Figure 2A) (Louie *et al*., 1988; Murphy *et al*., 1992; Ptitsyn, 1998). Besides, Vis-CD spectra, which give information about the coordination of the heme group (Figure 2B) were recorded. The peak observed around 410 nm corresponds to the B-band. The splitting of this band, which depends on the electronic perturbations of the heme group and its environment, indicated that octahedral coordination of the oxidized iron atom of both C*c* isoforms is maintained (Kelly *et al*., 2005; Hagarman *et al*., 2008).

**Figure 2.**
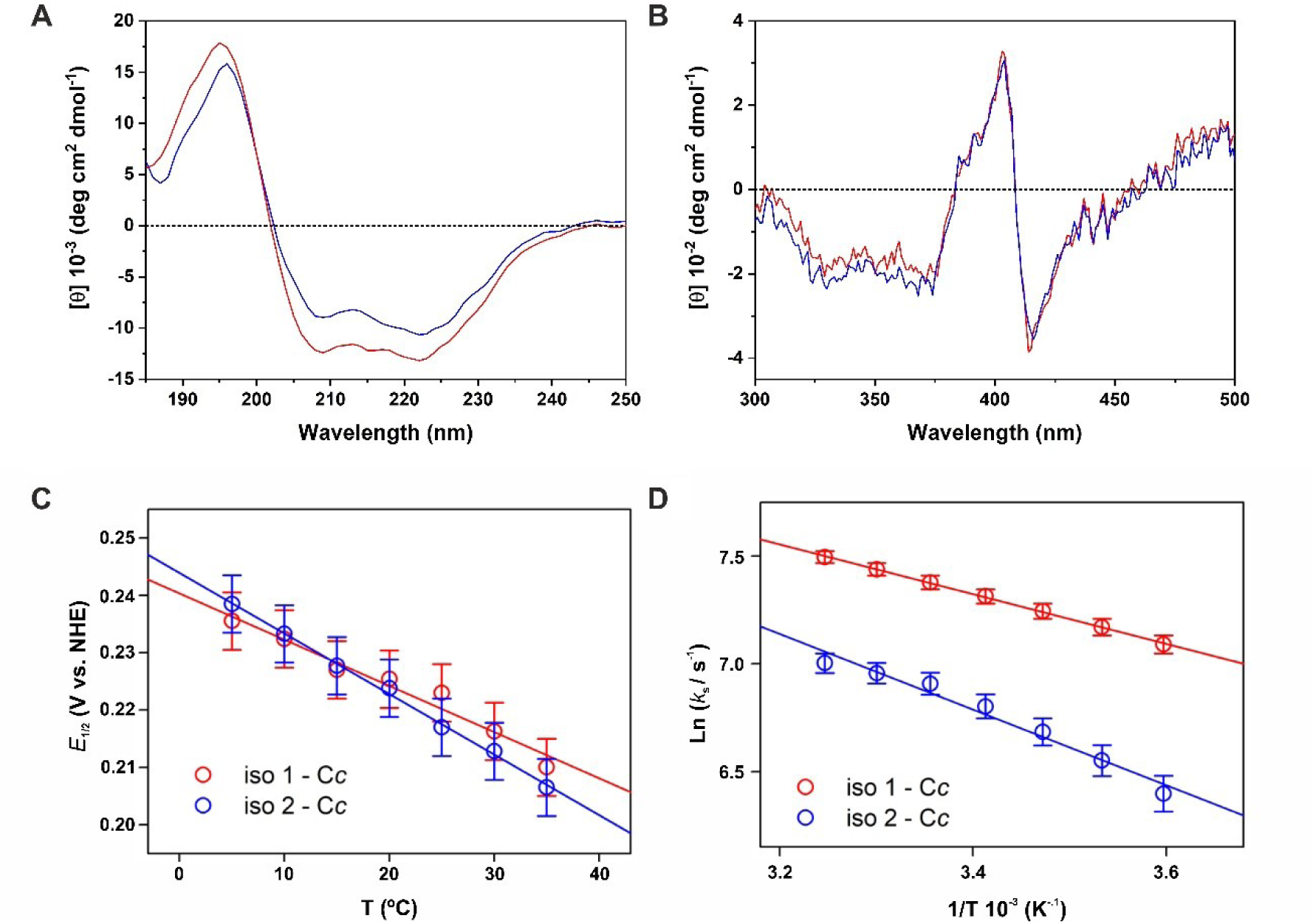
Circular dichroism spectra and electrochemical characterization of yeast cytochrome *c* isoforms. (**A**) Far-UV and (**B**) Vis CD spectra of oxidized iso-1 (red) and iso-2 (blue) C*c* species. (**C**) Variation of the midpoint potential *E*_1/2_, and (**D**) heterogeneous standard electron transfer rate constant *k*_*s*_ with the temperature for iso-1 (red symbols) and iso-2 (blue symbols) C*c* species immobilized onto a mixed-SAM-modified gold electrode and measured in 20 mM sodium phosphate buffer at pH 7.0. Solid lines correspond to the linear least-square fits of experimental data.

To assess the structural dissimilarity among the models of iso-1 and iso-2 C*c* (respectively, PDB accessions 3CX5 and 3CXH, Solmaz & Hunte, 2008), (Figure S1B in Annex A) we initially estimated the root-mean-square deviation (RMSD) values of both isoforms. A small value of 0.548 Å suggested that structural differences between yeast C*c* isoforms are negligible (Carugo & Pongor, 2001). To further explore differences in the molecular dynamics among both isoforms, 1 μs simulations of iso-1 and iso-2 were performed. Considering the tri-methylated state of naturally occurring yeast C*c*, unmodified and tri-methylated variants at Lys77 (in iso-1) or Lys81 (in iso-2) were included in the analysis as well. The backbone RMSD of the full-length constructs indicated that the iso-1 barely varies throughout the simulation, while iso-2 C*c* evolves from the initial structure during the first 100 ns of the simulation (Figure S3A in Annex A). Backbone root-mean-square fluctuations (RMSF) indicated that the 4 additional residues at the iso-2 C*c* N-terminus are notably flexible (Figure S3B in Annex A). In fact, restricting the RMSD analysis to the folded portion caused the RMSD values of all constructs to drop to ca. 1.5 Å (Figure S3C in Annex A), indicating that the slight deviation observed in RMSD values of the full-length protein is caused by the relaxation of the conformationally-constrained N-terminus in the crystal. Moreover, this region additionally causes a slight increase in the radius of gyration of the iso-2 compared to iso-1 (Figure S3D in Annex A). These extended flexible regions were predicted to not affect the region close to the heme crevice and the binding interface to electron transfer (Figure S3E in Annex A) and does not substantially impact the overall secondary structure of the protein (Figure S3F in Annex A). Taken together, these results evidence that the N-terminal region of iso-2 is conformationally more flexible than iso-1, which may have implications in the molecular recognition of electron transfer partners.

To gain information on the redox functionality of yeast C*c* isoforms in a physiological mimicking scenario, the proteins were adsorbed on a charged mixed monolayer of ω-mercaptocarboxylic acid and ω-hydroxy-n-alkanethiol modifying gold electrodes and the interfacial electron transfer of the proteins were characterized using cyclic voltammetry. Figure S4 (Appendix A) illustrates some typical voltammograms recorded at low scan rates of iso-1 and iso-2 C*c*, which display the shape characteristic of surface-confined redox species (Armstrong, 2002) consisting of well-defined reversible voltammetric waves that are associated with the heme Fe^3+^/Fe^2+^ redox conversion. The standard redox potentials of the two C*c* isoforms, as estimated from the midpoint potential (*E*_1/2_) of the anodic and cathodic voltammetric waves, were 0.223 V and 0.217 V vs. NHE for iso-1 C*c* and iso-2 C*c*, respectively. These *E*_1/2_ values were lower than the value of ∼ 290mV vs. NHE determined for both proteins in solution (de Groot *et al*., 2007), and are consistent with a relative stabilization of the protein ferric form following its electrostatic adsorption on a negatively charged thiol monolayer, previously observed for other mammalian, yeast, and bacterial cytochromes *c* (Fedurco, 2000; Todorovic *et al*., 2006; Oviedo-Rouco *et al*., 2022). Therefore, both isoforms increase their tendency to donate electrons upon electrostatic binding.

To determine thermodynamic and kinetic parameters accompanying their electron exchange with the electrode temperature-variable cyclic voltammetry was employed. The thermodynamic parameters 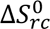 and 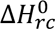 associated with the heme reduction were estimated from the temperature dependence of their *E*_1/2_ values based on (Taniguchi *et al*., 1980; Olloqui-Sariego *et al*., 2018; Olloqui-Sariego *et al*., 2019; Pérez-Mejías *et al*., 2020; Olloqui-Sariego *et al*., 2020; Márquez *et al*., 2021):

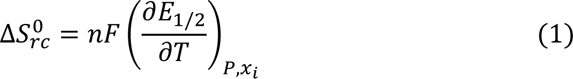

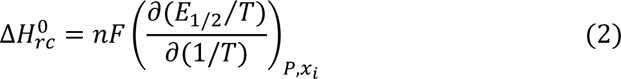

where *n* = 1 and *F* has its usual meaning.

The two C*c* isoforms display a linear decrease in *E*_1/2_ upon increasing the temperature (Figure 2C) which renders in negative 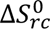 values (Table 1), according to equation 1. Specifically, entropic changes of -77.6 J K^-1^ mol^-1^ and -102 J K^-1^ mol^-1^ were estimated for iso-1 and iso-2 C*c*, respectively, which is consistent with a larger protein solvation change accompanying the redox conversion by iso-1. Analogously, the *E*_1/2_⁄*T* vs. *T*^−1^ plots (Figure S5 in Appendix A) also emerge negative tendencies, giving negative enthalpic changes values of -44.2 kJ mol^-1^ and -51.3 kJ mol^-1^ for the iso-1 and iso-2, respectively. These high negative enthalpies match with large differences in the stability of ligand binding interactions and hydrophobicity of the heme environment upon changing its oxidation state (Battistuzzi *et al*., 1999; Battistuzzi *et al*., 2004; di Rocco *et al*., 2021). Interestingly, the large absolute values determined for both isoforms involve a big enthalpy-entropy compensation which is consistent with an extensive rearrangement of its solvation shell after protein reduction.

**Table 1.** Thermodynamic and kinetic parameters of the Fe(III)/Fe(II) redox conversion of yeast cytochrome *c* isoforms.

|  | **iso-1 Cc** | **iso-2 Cc** |
| --- | --- | --- |
| $E_{1/2} / \text{mV}^a$ | 223 ± 5 | 217 ± 5 |
| $\Delta S_{rc}^0 / \text{J K}^{-1} \text{mol}^{-1}$ | -77.6 ± 6.0 | -102.0 ± 2.0 |
| $\Delta H_{rc}^0 / \text{kJ mol}^{-1}$ | -44.2 ± 1.9 | -51.3 ± 0.7 |
| $k_s / \text{s}^{-1}^a$ | 1600 ± 50 | 1000 ± 50 |
| $10^{-5} A / \text{s}^{-1}$ | 7.7 ± 0.6 | 34.5 ± 0.5 |
| $\Delta H_{ET}^\# / \text{kJ mol}^{-1}$ | 9.60 ± 0.12 | 14.60 ± 1.10 |

<sup>a</sup>Measured at 25 °C, pH 7.0

The kinetics of the electron transfer for each of the C*c* isoforms was also studied by evaluating the temperature-dependent behavior of the standard electron transfer rate constants (*k*_*s*_) for the Fe^3+^/Fe^2+^ redox conversion. *k*_*s*_ values were determined by fitting the variation of the voltammetric peak potential separation with the scan rate, employing a procedure based on Buttler-Volmer kinetic formalism with a transfer coefficient of 0.5 (Laviron, 1979). Symmetrical trumpet plots (Figure S6 in Appendix A) were obtained, expected for a well-behaved and quasi-reversible one-electron redox couple. The *k*_*s*_ value at 25 °C (Table 1) for iso-1 C*c* is slightly higher than the corresponding for iso-2

C*c*. To deepen into the molecular factors that govern the rate differences, the activation parameters of the electron transfer process of both C*c* isoforms were determined from the Arrhenius plots (Figure 2D). The intercept and the slope values of the linear Arrhenius plots provide pre-exponential factors (*A*) and activation enthalpies 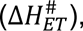, according to:

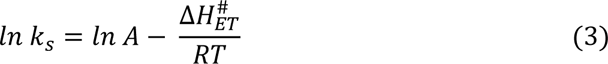

Interestingly, both 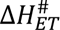 and A values of the isoforms were different (Table 1), being significantly higher for the iso-2. The lower 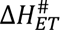 obtained for iso-1, can be justified by more restricted flexibility of the attached proteins and less participation of the solvent in the electron transfer process activation (Krishtalik, 2011), which also agrees with the estimated lower absolute values of the redox thermodynamics parameters of iso-2.

In addition, in terms of Marcus’ theory (Marcus, 1964; Marcus & Sutin, 1985), the pre-exponential relates with the donor-acceptor electronic coupling (*A* ∝ |*H*_*AB*_|^2^), which varies with the thickness and electronic conductivity of the intervening medium between electrode and redox center. Since the small differences in the amino acids sequence between two isoforms are not expected to induce dramatic changes in the electronic conductivity of the polypeptide matrix, the lower electronic coupling of iso-1 can be attributed to less productive protein orientations. Bearing that a lower electronic coupling decreases the interfacial electron transfer rate, but lower 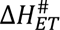 is consistent with a kinetically favored electron transfer, the higher *k*_*s*_ of the iso-1 arises from the higher contribution of the activation enthalpy. This result may suggest that the interaction of the isoforms with a charged interface involves different amino acids or orientations between redox centers.

### 3.2. *Saccharomyces cerevisiae* viability depends on the presence of cytochrome *c*

Five different yeast strains have been used: wild-type (WT); two deficient mutants in each of the C*c* isoforms (Δ*cyc1* and Δ*cyc7*, iso-1 and iso-2 C*c* deficient, respectively), and two complementary strains in each of the C*c* isoforms, which have been obtained from the mutant strains (*cyc1*^+^ and *cyc7*^+^, respectively).

Complementary strains were confirmed by PCR to amplify the *cyc1* and *cyc7* genes previously introduced in each of the complementary yeasts (Figure S7 in Appendix A). Bands correspond to DNA fragments between 300 and 400 base pairs. The electrophoretic mobility of the iso-2 C*c* band is slightly slower than the one corresponding to iso-1. This is consistent with the expected size for DNA fragments (*cyc1*: 330 bp and *cyc7*: 342 bp), thus corroborating the presence of the *cyc1* and *cyc7* genes in the respective complementary yeast strains.

The viability of the five yeast strains described earlier was tested by monitoring their growth patterns assay in YPD and YPG media (Figure S8 in Appendix A). In the YPD medium, yeast predominantly undergoes fermentative metabolism in the presence of oxygen, with glucose as a carbon source. To prevent glucose catabolism through respiratory pathways and the production of ethanol, cultures were collected at 24 h (Pfeiffer & Morley, 2014). On the other hand, in the YPG medium, yeast employs glycerol as a non-fermentable carbon source, imposing strictly respiratory metabolism and promoting the assembly of supercomplexes. Due to this metabolic restriction, cultures exhibited slower growth and were collected at 48 h to obtain a sufficient cellular mass for the assays (Figure S8 in Appendix A). All yeast strains grew in both media, allowing *in-mitochondria* studies.

Notably, it was attempted to obtain a double mutant strain deficient in both C*c* isoforms. This strain was not viable, even under a fermentable carbon source like glucose. The lethality of this double mutation for *S. cerevisiae* revealed that this yeast needs to have operational its ETC to survive. This finding is supported by other reports that describe how CIV does not assemble in the absence of C*c* or when it is not functional as an electron carrier (Bottorff *et al*., 1994; Pearce & Sherman, 1995; Vempati *et al*., 2009).

### 3.3. Respiratory supercomplexes assembly is modulated by cytochrome *c* isoforms

The assembly of respiratory supercomplexes mediated by C*c* isoforms has been analyzed using BN-PAGE based on solubilized mitochondria from the different strains grown in glucose (YPD medium, favoring fermentation) or glycerol (YPG medium, non-fermentable carbon sources, forcing respiratory metabolism) (Cruciat *et al*., 2000; Strogolova *et al*., 2012; Moreno-Beltrán *et al*., 2017; Timón-Gómez *et al*., 2020) (Figure 3). In this type of gel, proteins are in non-denaturing conditions, allowing the detection of interactions between them. Therefore, the band pattern provides information about the free and associated complexes into supercomplexes present in the five yeast strains grown in different media.

**Figure 3.**
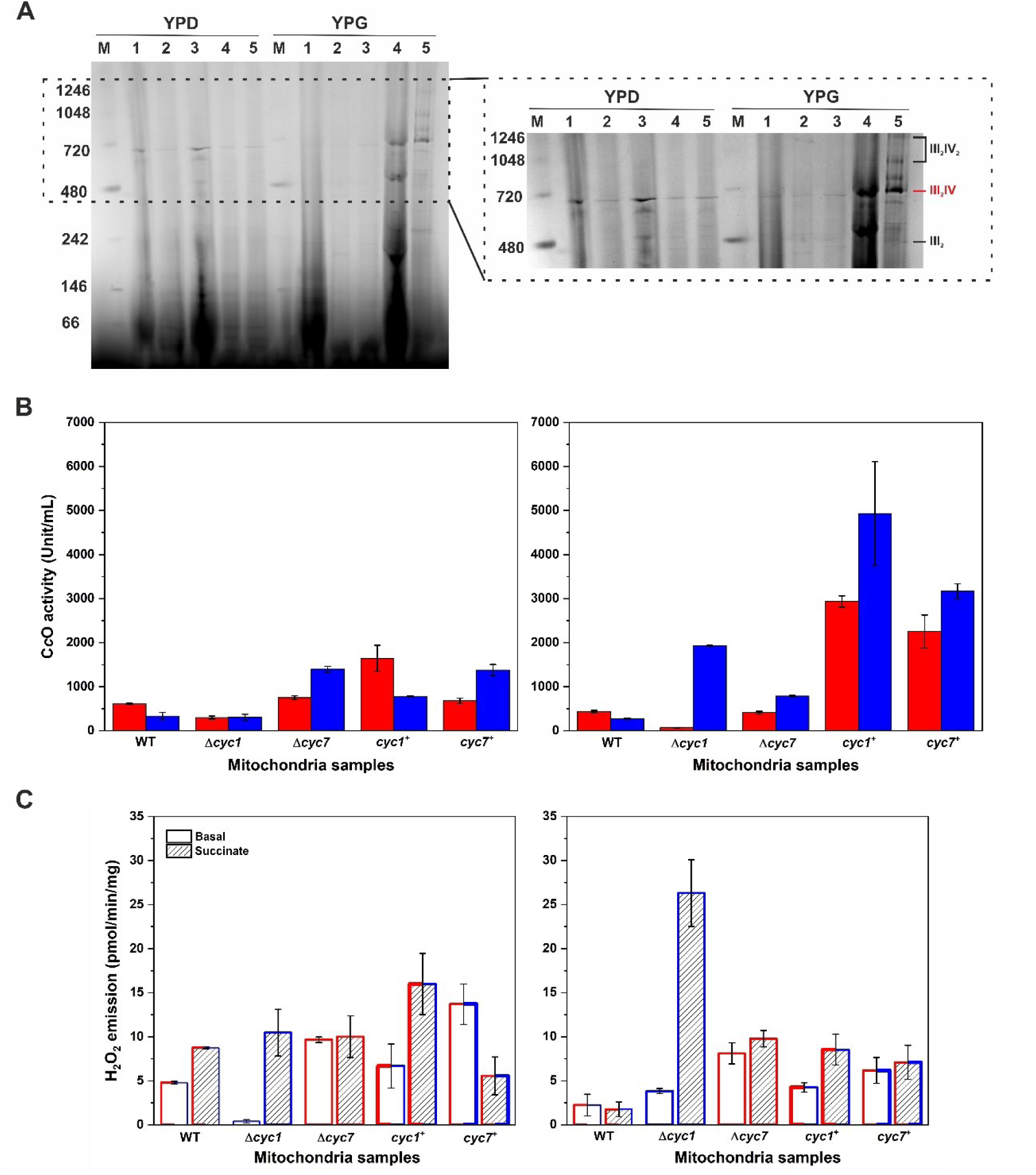
Supercomplexes formation and its impact on cytochrome *c* oxidase activity and reactive oxygen species production. (**A**) Analysis of respiratory supercomplex formation by BN-PAGE. Isolated mitochondria from strains: WT (1), Δ*cyc1* (2), Δ*cyc7* (3), *cyc1*^+^ (4) and *cyc7*^+^ (5), which have been grown in YPD (*left*) and YPG (*right*) media, and loaded into a BN-PAGE. Bands corresponding to III_2_IV supercomplex were submitted to tryptic digestion. M: molecular weight marker. (**B**) C*c*O activity *in mitochondria* from the following yeast strains: WT, Δ*cyc1*, Δ*cyc7*, *cyc1*^+^ and *cyc7*^+^ grown in YPD (*left*) and YPG (*right*) media and supplemented with iso-1 (red) and iso-2 (blue) C*c*. Activity values measured for each yeast strain upon adding exogenous iso-1 or iso-2 C*c*. (**D**) Reactive Oxygen Species (ROS) production in isolated mitochondria, as described in (**B**). *Basal*: ROS production before the addition of CII substrate. *Succinate*: ROS generation after the addition of succinate as NAD^+^-dependent substrate (CII). The colors of bars edges indicate the endogenous C*c* isoform present: iso-1 C*c* in red and iso-2 C*c* in blue. The edges of the bars are colored half red and half blue when both isoforms are present in WT, *cyc1*^+^ and *cyc7*^+^ strains. The edges of the bars are thicker to highlight the C*c* isoform that complements *cyc1*^+^ and *cyc7*^+^ strains.

In the YPD medium, a 720-kDa band was observed in all five yeast strains (Figure 3A), which corresponds to the assembly of supercomplex III_2_IV (Vukotic *et al*. 2012), being more intense in lanes of WT and *Δcyc2* yeast strains. Additionally, a 480-kDa band appeared in mitochondria isolated from Δ*cyc7* strains, corresponding to CIII dimer (III_2_) (Timón-Gómez *et al*., 2020).

In YPG medium, all strains showed the previously mentioned 720-kDa band, although it was particularly intense and prominent in the complementary yeast strains *cyc1*^+^ and *cyc7*^+^. Moreover, these strains displayed additional bands: while the 480-kDa band was clearly evident in *cyc1*^+^ strains, bands upper the one at 720 kDa corresponding to the supercomplex III_2_IV_2_, along with assembly factors co-migrating with this supercomplex, were present in *cyc7*^+^ strains (Timón-Gómez *et al*., 2020).

We analyzed and identified the protein composition of the 720-kDa band across all lanes, and could successfully identify the different C*c* isoforms —in agreement with each strain’s expression based on its genotype (Table 2; and Appendix C along with Excel files,). These findings showed that C*c* forms a stable interaction with III_2_IV supercomplex, and might function as a modulator for the assembly process.

**Table 2.** Identified proteins by tryptic digestion analyses.

| **Lane** | **Protein** |
| --- | --- |
| YPD-1 (WT) | Cc<sub>1</sub> (CIII)<br>Rieske subunit (CIII)<br>COX5A (CIV)<br>COX5B (CIV)<br>**iso-1 Cc**<br>**iso-2 Cc** |
| YPD-2 ($\Delta cyc1$) | Cc<sub>1</sub> (CIII)<br>Rieske subunit (CIII)<br>COX5A (CIV)<br>ATP synthase subunit 4<br>**iso-2 Cc** |
| YPD-3 ($\Delta cyc2$) | COX5A |
| YPD-4 (*cyc1*<sup>+</sup>) | - |
| YPD-4 (*cyc2*<sup>+</sup>) | - |
| YPG-1 (WT) | Cc<sub>1</sub> (CIII)<br>Rieske subunit (CIII)<br>**iso-1 Cc**<br>**iso-2 Cc** |
| YPG-2 ($\Delta cyc1$) | - |
| YPG-3 ($\Delta cyc2$) | - |
| YPG-4 (*cyc1*<sup>+</sup>) | COX5A (CIV)<br>ATP synthase subunit alpha |
| YPG-5 (*cyc2*<sup>+</sup>) | Cytochrome *bc*<sub>1</sub> complex subunit 1 (CIII)<br>Cytochrome *bc*<sub>1</sub> complex subunit 2 (CIII)<br>Cc<sub>1</sub> (CIII)<br>Rieske subunit (CIII)<br>Cytochrome c oxidase subunit 2<br>Cytochrome c oxidase polypeptide VIII (CIII)<br>COX5A (CIV)<br>COX5B (CIV)<br>ATP synthase subunit alpha<br>ATP synthase subunit beta<br>ATP synthase subunit gamma<br>ATP synthase subunit epsilon<br>ATP synthase subunit f<br>ATP synthase subunit J<br>ATP synthase subunit g<br>ATP synthase subunit 4<br>ATP synthase subunit 5<br>**iso-1 Cc**<br>**iso-2 Cc** |

Taken together, despite the prevalence of fermentative metabolism in the YPD medium, yeasts can assemble supercomplexes and keep the ETC operational to ensure the integrity of the inner mitochondrial membrane (IMM) and, consequently, maintain physiological ATP and oxidative levels, as well as the correct functioning of signaling processes to avoid apoptosis (Hong *et al*., 2022). Thus, the functions of ETC components may extend beyond energy metabolism, which likely explains why the yeast double knock-out strain (ΔΔ*cyc1cyc7*) was inviable (Zhang *et al*., 2015; Sedlackova & Korolchuk, 2019). Notably, the assembly of supercomplexes in the YPG medium is dependent on the presence or absence of C*c* isoforms. Whereas iso-1 C*c* is essential for the assembly of respiratory III_2_IV supercomplex, iso-2 C*c* is relevant to form not only III_2_IV but also larger, highly-ordered ones such as III_2_IV_2_ supercomplex. In summary, both iso-1 and iso-2 hemeproteins act as modulators of supercomplexes assembly.

### 3.4. Isoform 2 of cytochrome *c* is more efficient in transferring electrons towards cytochrome *c* oxidase

The ability of the different C*c* isoforms as electron carriers was analyzed by measuring their C*c*O activity. In the presence of isolated CIV, the C*c*O activity was higher upon the addition of exogenous iso-2 C*c* (Figure S9 in Appendix A).

In mitochondria isolated from yeast Δ*cyc1* strains and grown in YPD medium, C*c*O activity decreases with respect to mitochondria from the WT strain upon adding exogenous iso-1 C*c* (Figure 3C, red bars). The addition of iso-2 C*c*, however, keeps the enzymatic activity mainly unchanged (Figure 3B, blue bars). A plausible explanation is the scarce supercomplex formation in Δ*cyc1* yeasts under the YPD medium (Figure 3A). However, C*c*O activity increases when both C*c* isoforms are added to Δ*cyc7* strains, in line with the presence of supercomplexes (Figures 3A and 3B). *Cyc1*+ strains show activity levels larger than those from WT strains, regardless of the isoform incorporated. Interestingly, the enzymatic levels of *cyc7*+ strains closely resemble those observed for Δ*cyc7* strains. While supercomplex bands are present in the absence of *cyc7* (Δ*cyc7* strains), iso-2 C*c* overexpression in *cyc7*+ strains is responsible for a higher turnover rate of the hemeprotein donating electrons to CIV.

Mitochondria isolated from cells grown in YPG medium showed increased C*c*O activity for those cultured in YPD medium in such mitochondria with high-ordered complex structures and/or supplemented with iso-2 C*c* (Figure 3B). Further differences in enzymatic activity observed between the two yeasts complemented with iso-1 and iso-2 C*c* might be explained considering different adsorption onto the negatively charged thiol monolayers, potentially resulting in less efficient orientations for iso-1 C*c*.

ROS production was also analyzed because the changes in the electron flow through the ETC due to the assembly of respiratory supercomplexes can affect the hyperpolarization of the IMM and ROS production (Starkov & Fiskum, 2003). In the yeast *S. cerevisiae*, ROS generation primarily occurs at CIII since this organism lacks CI, but ROS production also takes place at CIV (De Vries & Grivell, 1988; Herrero *et al*., 2008).

To stimulate ROS production, succinate was used as a substrate for CII. It is important to remark that no exogenous species of C*c* were added in these assays, so the observed ROS production is due to the formation of supercomplexes and the endogenous isoform present in each strain (Figure 3C).

In the YPD medium (Figure 3C, *left*), mitochondria isolated from WT strains show a progressive increase in ROS production when the ETC is triggered. A similar situation is described for mitochondria from Δ*cyc1* strain, although it exhibits a very low basal level that increases when succinate is added. In Δ*cyc7* strain, the presence of supercomplexes helps to maintain similar ROS levels throughout the assay, although the basal ROS level is larger in comparison with other strains. For complementary yeasts, the behavior is opposite between *cyc1+* and *cyc7+* strains. H_2_O_2_ production increased upon succinate addition in *cyc1+* strain, while the constitutive expression of iso-2 C*c* in *cyc7*+ strain resulted in a reduction of ROS.

In YPG medium, the presence of supercomplexes in WT strain maintains ROS levels low after succinate addition. In Δ*cyc1* strain —where supercomplex formation is not observed— once the ETC is triggered by succinate, ROS production increases significantly. However, supercomplex assembly in the Δ*cyc7* strain keeps ROS production levels lower. The overexpression of iso-1 C*c* in *cyc1*^+^ strain triggered supercomplex assembly and kept ROS low when succinate was added. In the case of *cyc7*^+^ strain, ROS levels remain low in the presence of succinate. This could be explained based on the supercomplex bands above 720 kDa observed in this strain and/or the higher efficiency of iso-2 C*c* in electron transfer.

## 4. Conclusions

The two yeast C*c* isoforms maintain the overall secondary structure and the heme coordination, although iso-2 C*c* donates electrons more efficiently towards C*c*O. Under restricted respiratory conditions, high-ordered supercomplexes (III_2_IV and III_2_IV_2_) are assembled in complementary yeast strains (*cyc1*^+^ and *cyc7*^+^), underscoring the critical role of both iso-1 and iso-2 C*c* in these adducts. Noteworthy, the III_2_IV and III_2_IV_2_ supercomplexes, together with iso-2 C*c*, lead to a productive and efficient mitochondrial electron flow that keeps ROS levels low. Since there are no significant variations in the residues participating in the binding sites among the two isoforms, differences in the electron carrier ability among C*c* isoforms in the supercomplex context could be attributed to the distance and relative orientation between the redox centers, similarly as happens in other redox complexes (Navarro *et al*., 1995; Sun *et al*., 1999; Díaz-Moreno *et al*., 2005; Cruz-Gallardo *et al*., 2012).

The first, important consequence of respiratory III_2_IV and III_2_IV_2_ supercomplex arrangement is shortening the distance that C*c* has to go through to transfer electrons between its redox partners, along with restricted the search for an efficient electron transfer from 3D to 2D (Table S3 in Appendix A) (Berndtsson *et al*., 2020; Pérez-Mejías *et al*., 2022). This explains a higher C*c*O activity followed by a lower ROS production (Guan *et al*., 2022). In the supercomplexes context, the distance involved in the electron transfer between C*c* and its redox partners was shorter in the case of iso-2 C*c*, consisting of the higher efficiency of this isoform (Table S3 in Appendix A).

Altogether, these findings suggest that the C*c*O-driven oxidation rate is regulated by i) the formation and type of supercomplexes; ii) the isoform of C*c*, so as to keep apoptotic-inducing mitochondrial ROS at a low level. This implies a regulatory mechanism through the respiratory supercomplexes assembly modulated by C*c* to achieve a cellular response to oxidative stress, hypoxia, or other respiratory or signalling needs. Thus, C*c* emerges as a new supercomplex assembly factor under hypoxia, amplifying its cell role as a signaller beyond electron carrier and programmed cell death inductor (Ow YP *et al*., 2008; Martínez-Fábregas J et al, 2014; González-Arzola K *et al*., 2019).

## Supporting information

Tryptic Digestion Analyses Of Cytochrome c Isoforms

Tryptic Digestion Analyses of BN-PAGE

## Abbreviations

*cyc1^+^*: cytochrome *c* isoform 1 complementary yeast strain
*cyc7^+^*: cytochrome *c* isoform 2 complementary yeast strain
Δ*cyc1*: cytochrome c isoform 1 deficient yeast
Δ*cyc7*: cytochrome *c* isoform 2 deficient yeast
iso-1: isoform 1
iso-2: isoform 2
MES: 2-(N-morpholino)ethanesulfonic acid, 4-morpholineethanesulfonic acid
MOA: 8 mercaptooctanoic acid
MOOL: 8-mercapto-1-octanol
NHE: normal hydrogen electrode
Q: quinone
SAM: self-assembled monolayers
YPD: yeast extract peptone dextrose
YPG: yeast extract peptone glycerol

## Declaration of competing interests

The authors declare no conflicts of interest.

## Data availability

Data support the findings of this study are available from the corresponding authors upon request.

## Acknowledgments

We acknowledge Prof. Carlos Santos Ocaña (Andalusian Center for Developmental Biology, CABD) for the Δ*cyc1* strain. We also acknowledge the facility and scientific and technical assistance of the Biomolecular Mass Spectrometry Service of Pablo de Olavide University.

## Funding sources

This work was supported by the Spanish Government (PID2021-126663NB-I00 funded by MCIN/AEI/ 10.13039/501100011033 and by ERDF A way of making Europe), European Regional Development Fund (FEDER), Andalusian Government (BIO-198, US/JUNTA/FEDER) and the Ramón Areces Foundation (2021-2024 to I.D.-M.).

## Appendices. Supplementary data

Appendix A. Supplementary Figures And Tables Appendix C. Tryptic Digestion Analyses of BN-PAGE

## Notes

### Competing Interest Statement

The authors have declared no competing interest.

