## Supplementary material for "Unveiling the role of yeast cytochrome *c* isoforms in the assembly of mitochondrial supercomplexes and the control of respiratory chain rate": Tryptic Digestion Analyses Of Cytochrome c Isoforms

#### APPENDIX B. TRYPTIC DIGESTION ANALYSES OF CYTOCHROME *c* ISOFORMS

Alejandra Guerra-Castellano[a,*], Manuel Aneas[a], Joaquín Tamargo-Azpilicueta[a], Inmaculada Márquez[b], José Luis Olloqui-Sariego[b], Juan José Calvente[b], Miguel A. De la Rosa[a], Irene Díaz-Moreno[a,*]

[a]Instituto de Investigaciones Químicas, Centro de Investigaciones Científicas Isla de la Cartuja, Universidad de Sevilla - CSIC. Avda. Americo Vespucio 49, 41092 Sevilla, Spain.
[b]Departamento de Química Física, Universidad de Sevilla, Profesor García González 1, 41012 Sevilla, Spain.

#### Spectrum Analysis Report

### iso-1 Cc

| | | | | | | | |
|---|---|---|---|---|---|---|---|
| Sequence Name: | | Formula: | | Parentmass: | | Mass Error: | |
| MH+ (mono): | 1.008 | MH+ (avg): | 1.008 | Threshold (a.i.): | 0.000 | Tolerance (Da): | 0.500 |
| Number of Peaks: | 22 | Above Threshold: | | Assigned Peaks: | | Not assigned Peaks: | |

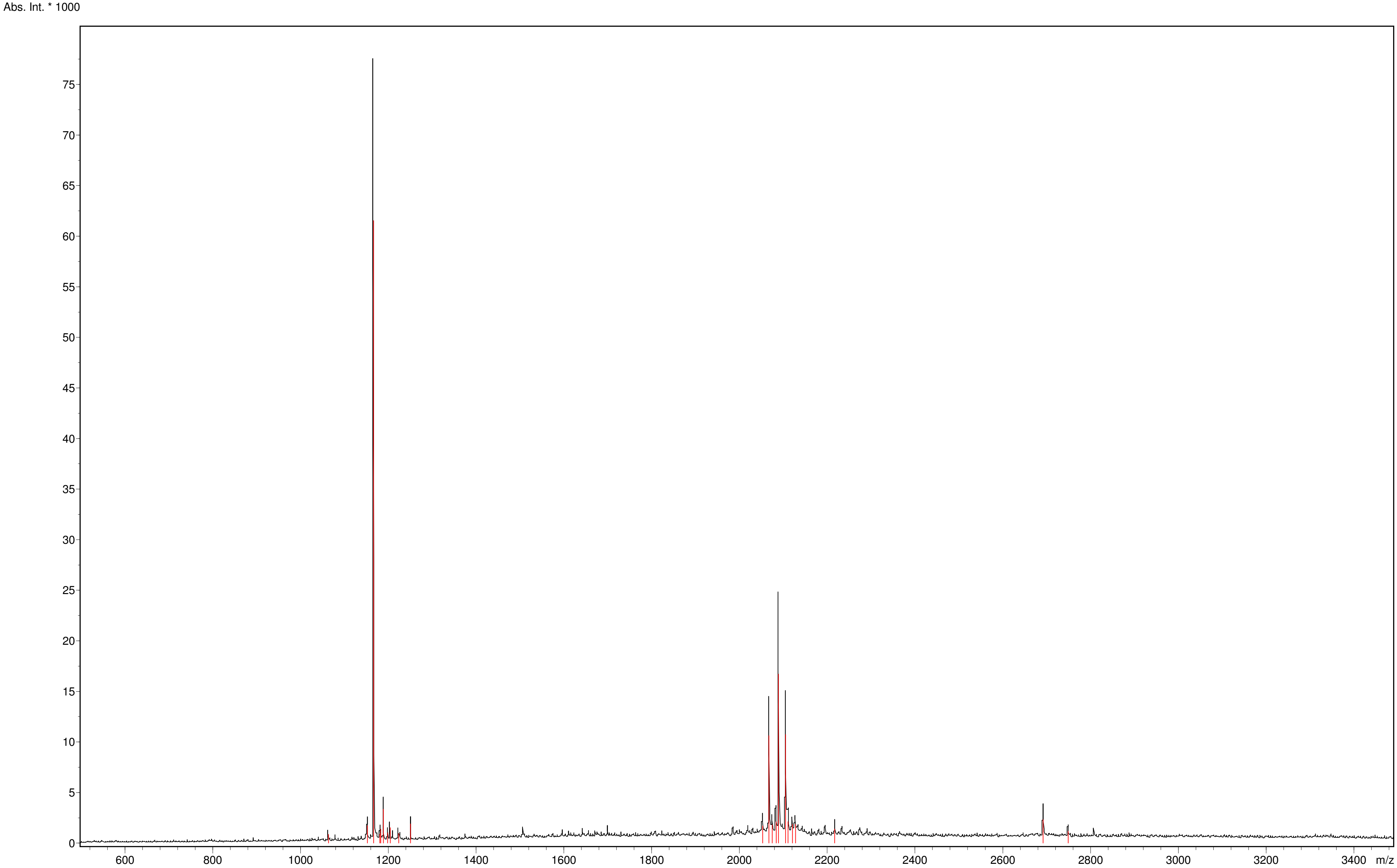



Copyright © 1999-2008 Bruker Daltonik GmbH

#### Spectrum Analysis Report
### iso-1 Cc

**Sequence data:**

Intensity Coverage: 49.0 % (63436 cnts)
Sequence Coverage MS/MS: 0.0%
Sequence Coverage MS: 19.3%
pI (isoelectric point): 10.1

```
        10         20         30         40         50         60         70         80         90        100        110
MTEFKAGSAK KGATLFKTRC LQCHTVEKGG PHKVGPNLHG IFGRHSGQAE GYSYTDANIK KNVLWDENNM SEYLTNPKKY IPGTKMAFGG LKKEKDRNDL ITYLKKACE
```

**Display Parameter:**

MH+ (mono): 1.008 MH+ (avg): 1.008 Threshold (a.i.): 0.000 Tolerance (Da): 0.500
Number of Peaks: 22

**Peaklist:**

| Peak | Mass | Intensity | Peak | Mass | Intensity | Peak | Mass | Intensity | Peak | Mass | Intensity | Peak | Mass | Intensity |
|---|---|---|---|---|---|---|---|---|---|---|---|---|---|---|
| 1 | 1063.753 | 872.381 | 2 | 1152.606 | 1716.103 | 3 | 1166.677 | 61534.508 | 4 | 1180.605 | 914.145 | 5 | 1182.651 | 1383.747 |
| 6 | 1188.645 | 3341.283 | 7 | 1198.657 | 1013.491 | 8 | 1204.632 | 1533.415 | 9 | 1223.663 | 958.483 | 10 | 1250.686 | 1901.593 |
| 11 | 2053.027 | 1711.415 | 12 | 2067.055 | 10625.056 | 13 | 2074.752 | 1812.163 | 14 | 2083.954 | 2110.268 | 15 | 2089.005 | 16701.365 |
| 16 | 2104.988 | 10727.034 | 17 | 2110.935 | 2113.130 | 18 | 2120.969 | 1852.827 | 19 | 2127.925 | 1865.367 | 20 | 2217.139 | 1391.165 |
| 21 | 2692.091 | 2327.563 | 22 | 2749.085 | 1168.750 | | | | | | | | | |


Copyright © 1999-2008 Bruker Daltonik GmbH

**Spectrum Analysis Report**

#### iso-2 Cc

| | | | | | | | |
|---|---|---|---|---|---|---|---|
| Sequence Name: | | Formula: | | Parentmass: | | Mass Error: | |
| MH+ (mono): | 1.008 | MH+ (avg): | 1.008 | Threshold (a.i.): | 0.000 | Tolerance (Da): | 0.500 |
| Number of Peaks: | 50 | Above Threshold: | | Assigned Peaks: | | Not assigned Peaks: | |

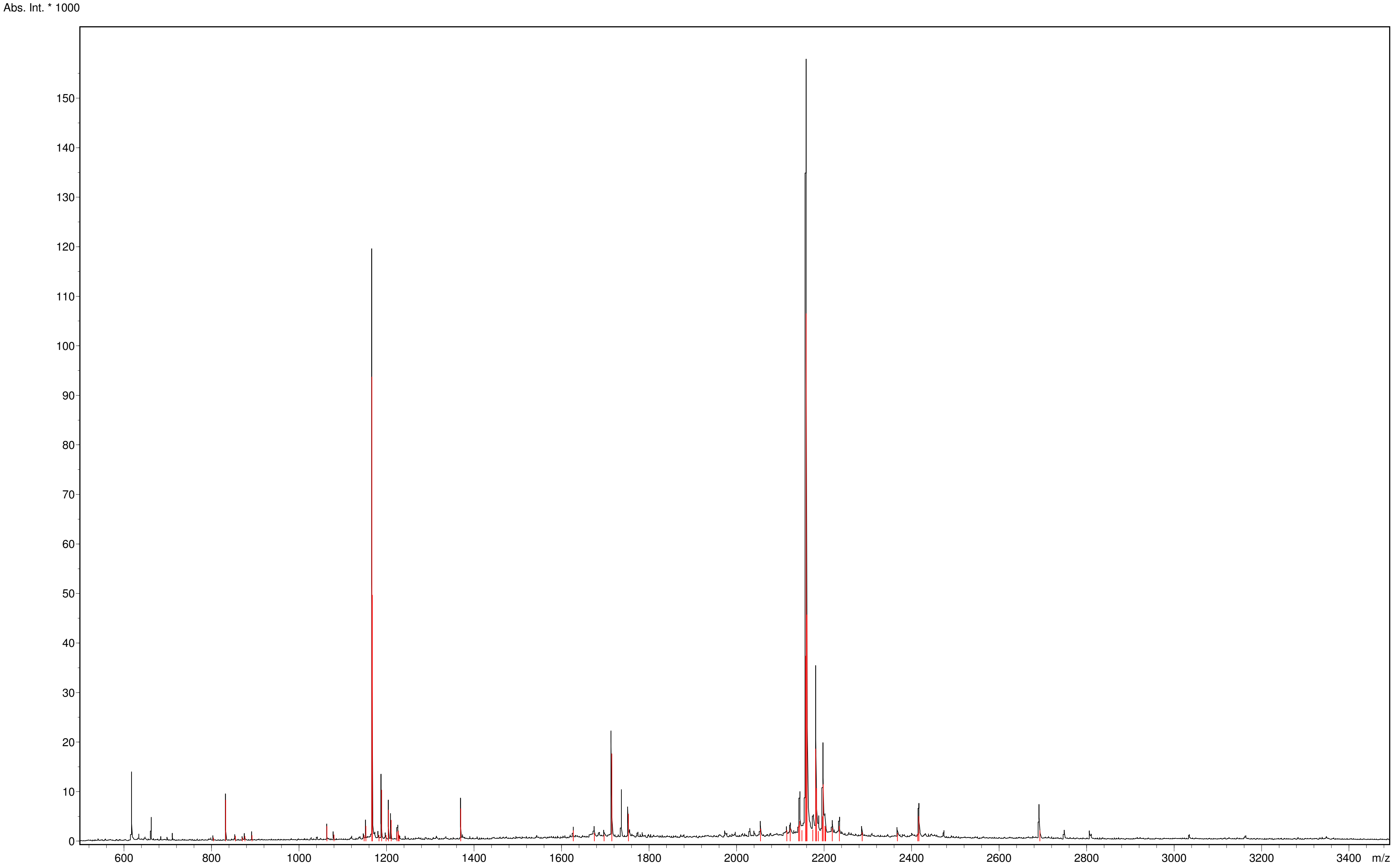



Copyright © 1999-2008 Bruker Daltonik GmbH

#### Spectrum Analysis Report
#### iso-2 Cc

**Sequence data:**

| | | | |
|---|---|---|---|
| Intensity Coverage: | 23.7 % (119988 cnts) | Sequence Coverage MS: | 33.6% |
| Sequence Coverage MS/MS: | 0.0% | pI (isoelectric point): | 10.2 |

| 10 | 20 | 30 | 40 | 50 | 60 | 70 | 80 | 90 | 100 | 110 |
|---|---|---|---|---|---|---|---|---|---|---|
| MAKESTGFKP | GSAKKGATLF | KTRCQQCHTI | EEGGPNKVGP | NLHGIFGRHS | GQVKGYSYTD | ANINKNVKWD | EDSMSEYLTN | PKKYIPGTKM | AFAGLKKEKD | RNDLITYMTK |
| 120 | | | | | | | | | | |
| AAK | | | | | | | | | | |

**Display Parameter:**

| | | | | | | | |
|---|---|---|---|---|---|---|---|
| MH+ (mono): | 1.008 | MH+ (avg): | 1.008 | Threshold (a.i.): | 0.000 | Tolerance (Da): | 0.500 |
| Number of Peaks: | 50 | | | | | | |

**Peaklist:**

| Peak | Mass | Intensity | Peak | Mass | Intensity | Peak | Mass | Intensity | Peak | Mass | Intensity | Peak | Mass | Intensity |
|---|---|---|---|---|---|---|---|---|---|---|---|---|---|---|
| 1 | 804.324 | 633.705 | 2 | 832.363 | 8288.950 | 3 | 854.326 | 993.274 | 4 | 870.311 | 629.878 | 5 | 876.294 | 941.459 |
| 6 | 892.279 | 1145.816 | 7 | 1063.766 | 2953.578 | 8 | 1079.679 | 1266.398 | 9 | 1148.628 | 1123.359 | 10 | 1152.600 | 3025.819 |
| 11 | 1166.649 | 93692.310 | 12 | 1167.631 | 49670.271 | 13 | 1182.630 | 1233.521 | 14 | 1188.620 | 10317.432 | 15 | 1198.635 | 1198.744 |
| 16 | 1204.597 | 6286.266 | 17 | 1210.596 | 4234.755 | 18 | 1223.653 | 1847.938 | 19 | 1226.565 | 2090.305 | 20 | 1228.535 | 1209.715 |
| 21 | 1369.679 | 6539.363 | 22 | 1626.819 | 2115.901 | 23 | 1674.837 | 1611.043 | 24 | 1696.924 | 1288.782 | 25 | 1714.708 | 17640.022 |
| 26 | 1752.684 | 5470.199 | 27 | 2054.453 | 2488.870 | 28 | 2114.862 | 2185.033 | 29 | 2122.514 | 2545.258 | 30 | 2141.797 | 3157.077 |
| 31 | 2144.502 | 4018.877 | 32 | 2149.218 | 2170.523 | 33 | 2157.766 | 37346.554 | 34 | 2158.901 | 106500.534 | 35 | 2160.794 | 45695.683 |
| 36 | 2174.439 | 2290.245 | 37 | 2180.864 | 18644.530 | 38 | 2182.763 | 10648.072 | 39 | 2186.970 | 1992.380 | 40 | 2195.796 | 4922.808 |
| 41 | 2197.800 | 11414.128 | 42 | 2201.801 | 3051.842 | 43 | 2203.770 | 2871.078 | 44 | 2218.839 | 2602.189 | 45 | 2234.836 | 3017.247 |
| 46 | 2286.935 | 1715.678 | 47 | 2367.885 | 1598.009 | 48 | 2414.090 | 1563.582 | 49 | 2417.003 | 5000.695 | 50 | 2693.240 | 2103.986 |


Copyright © 1999-2008 Bruker Daltonik GmbH
