## Supplementary material for "Unveiling the role of yeast cytochrome *c* isoforms in the assembly of mitochondrial supercomplexes and the control of respiratory chain rate": Tryptic Digestion Analyses of BN-PAGE

**APPENDIX C. TRYPTIC DIGESTION ANALYSES OF BN-PAGE**

Alejandra Guerra-Castellano[a,*], Manuel Aneas[a], Joaquín Tamargo-Azpilicueta[a], Inmaculada Márquez[b], José Luis Olloqui-Sariego[b], Juan José Calvente[b], Miguel A. De la Rosa[a], Irene Díaz-Moreno[a,*]

[a]Instituto de Investigaciones Químicas, Centro de Investigaciones Científicas Isla de la Cartuja, Universidad de Sevilla - CSIC. Avda. Americo Vespucio 49, 41092 Sevilla, Spain.
[b]Departamento de Química Física, Universidad de Sevilla, Profesor García González 1, 41012 Sevilla, Spain.

**Unveiling the role of yeast cytochrome *c* isoforms in the assembly of mitochondrial supercomplexes and the control of respiratory chain rate**

#### ANNEX C. TRYPTIC DIGESTION ANALYSES OF BN-PAGE

### YPD-1

#### Spectrum Analysis Report

### Cytohrome c1

| | | | | | | | |
|---|---|---|---|---|---|---|---|
| Sequence Name: | | Formula: | | Parentmass: | | Mass Error: | |
| MH+ (mono): | 1.008 | MH+ (avg): | 1.008 | Threshold (a.i.): | 0.000 | Tolerance (Da): | 0.500 |
| Number of Peaks: | 74 | Above Threshold: | | Assigned Peaks: | | Not assigned Peaks: | |

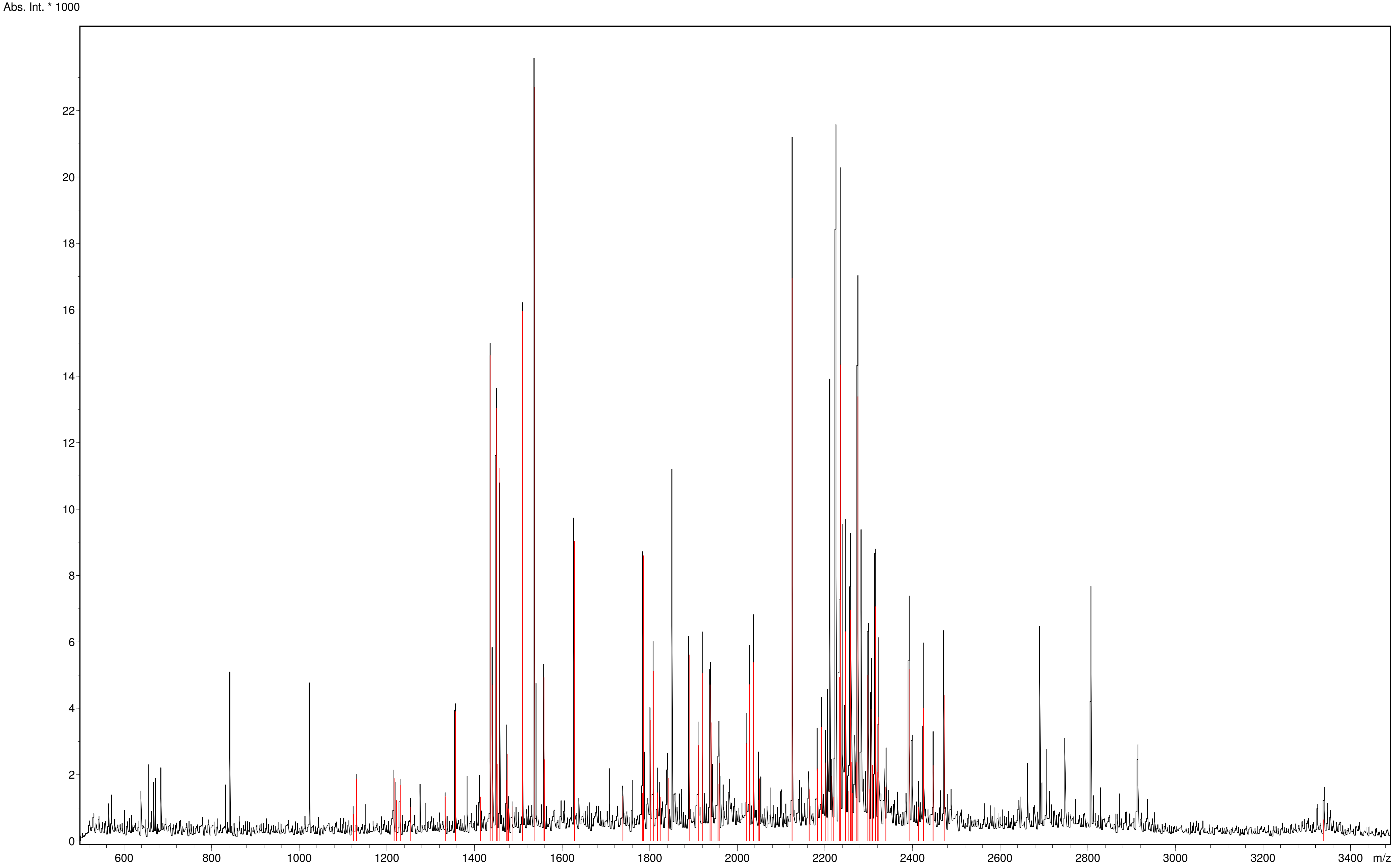



Copyright © 1999-2008 Bruker Daltonik GmbH

#### Spectrum Analysis Report

### Cytochrome c1

**Sequence data:**

| | | | |
|---|---|---|---|
| Intensity Coverage: | 2.0 % (6304 cnts) | Sequence Coverage MS: | 5.8% |
| Sequence Coverage MS/MS: | 0.0% | pI (isoelectric point): | 9.0 |

| 10 | 20 | 30 | 40 | 50 | 60 | 70 | 80 | 90 | 100 | 110 | 120 |
|---|---|---|---|---|---|---|---|---|---|---|---|
| MFSNLSKRWA | QRTLSKSFYS | TATGAASKSG | KLTQKLVTAG | VAAAGITAST | LLYADSLTAE | AMTAAEHGLH | APAYAWSHNG | PFETFDHASI | RRGYQVYREV | CAACHSLDRV | AWRTLVGVSH |
| **130** | **140** | **150** | **160** | **170** | **180** | **190** | **200** | **210** | **220** | **230** | **240** |
| TNEEVRNMAE | EFEYDDEPDE | QGNPKKRPGK | LSDYIPGPYP | NEQAARAANQ | GALPPDLSLI | VKARHGGCDY | IFSLLTGYPD | EPPAGVALPP | GSNYNPYFPG | GSIAMARVLF | DDMVEYEDGT |
| **250** | **260** | **270** | **280** | **290** | **300** | **310** | | | | | |
| PATTSQMAKD | VTTFLNWCAE | PEHDERKRLG | LKTVIILSSL | YLLSIWVKKF | KWAGIKTRKF | VFNPPKPRK | | | | | |

**Display Parameter:**

| | | | | | | | |
|---|---|---|---|---|---|---|---|
| MH+ (mono): | 1.008 | MH+ (avg): | 1.008 | Threshold (a.i.): | 0.000 | Tolerance (Da): | 0.500 |
| Number of Peaks: | 74 | | | | | | |

**Peaklist:**

| Peak | Mass | Intensity | Peak | Mass | Intensity | Peak | Mass | Intensity | Peak | Mass | Intensity | Peak | Mass | Intensity |
|---|---|---|---|---|---|---|---|---|---|---|---|---|---|---|
| 1 | 1123.583 | 808.568 | 2 | 1130.622 | 1869.448 | 3 | 1216.596 | 1883.277 | 4 | 1221.606 | 1226.315 | 5 | 1230.633 | 1677.819 |
| 6 | 1254.531 | 1032.225 | 7 | 1333.672 | 1337.183 | 8 | 1356.622 | 3905.019 | 9 | 1413.702 | 1328.696 | 10 | 1435.724 | 14621.908 |
| 11 | 1441.744 | 4709.528 | 12 | 1449.741 | 13033.796 | 13 | 1451.725 | 2323.019 | 14 | 1457.727 | 11235.150 | 15 | 1472.716 | 1826.124 |
| 16 | 1474.723 | 2623.582 | 17 | 1477.723 | 971.750 | 18 | 1485.749 | 1046.601 | 19 | 1509.671 | 15966.638 | 20 | 1537.725 | 22703.080 |
| 21 | 1558.823 | 4932.817 | 22 | 1559.714 | 2454.025 | 23 | 1627.815 | 9030.708 | 24 | 1738.836 | 1349.505 | 25 | 1783.819 | 1440.397 |
| 26 | 1785.965 | 8593.898 | 27 | 1800.915 | 3643.763 | 28 | 1807.949 | 5120.049 | 29 | 1817.848 | 1461.242 | 30 | 1823.906 | 1323.379 |
| 31 | 1841.902 | 1891.399 | 32 | 1890.059 | 5611.720 | 33 | 1912.043 | 2878.011 | 34 | 1919.942 | 5041.679 | 35 | 1937.979 | 4709.868 |
| 36 | 1941.934 | 3570.011 | 37 | 1955.951 | 1710.427 | 38 | 1959.966 | 2346.213 | 39 | 2021.032 | 2934.943 | 40 | 2027.950 | 4711.074 |
| 41 | 2037.024 | 5380.085 | 42 | 2048.923 | 1236.841 | 43 | 2051.005 | 1878.804 | 44 | 2125.105 | 16947.955 | 45 | 2164.078 | 1550.771 |
| 46 | 2183.020 | 2179.902 | 47 | 2192.118 | 3430.623 | 48 | 2202.102 | 2358.338 | 49 | 2207.097 | 2693.954 | 50 | 2214.134 | 1944.392 |
| 51 | 2220.090 | 1328.941 | 52 | 2233.079 | 4933.313 | 53 | 2236.103 | 14340.864 | 54 | 2247.118 | 6304.303 | 55 | 2253.096 | 1507.950 |
| 56 | 2258.092 | 6961.212 | 57 | 2261.169 | 2382.070 | 58 | 2262.661 | 1441.921 | 59 | 2273.062 | 2367.695 | 60 | 2275.191 | 13389.789 |
| 61 | 2298.170 | 5004.297 | 62 | 2301.153 | 1564.191 | 63 | 2305.138 | 3968.486 | 64 | 2308.101 | 2283.139 | 65 | 2315.167 | 7059.208 |
| 66 | 2320.158 | 2085.198 | 67 | 2323.096 | 3727.940 | 68 | 2339.098 | 1628.892 | 69 | 2392.209 | 5182.606 | 70 | 2414.193 | 1030.947 |
| 71 | 2425.200 | 4004.982 | 72 | 2447.172 | 2280.361 | 73 | 2472.270 | 4390.654 | 74 | 3338.700 | 645.613 | | | |


Copyright © 1999-2008 Bruker Daltonik GmbH

#### Spectrum Analysis Report
### Rieske subunit

| | | | | | | | |
|---|---|---|---|---|---|---|---|
| Sequence Name: | | Formula: | | Parentmass: | | Mass Error: | |
| MH+ (mono): | 1.008 | MH+ (avg): | 1.008 | Threshold (a.i.): | 0.000 | Tolerance (Da): | 0.500 |
| Number of Peaks: | 74 | Above Threshold: | | Assigned Peaks: | | Not assigned Peaks: | |

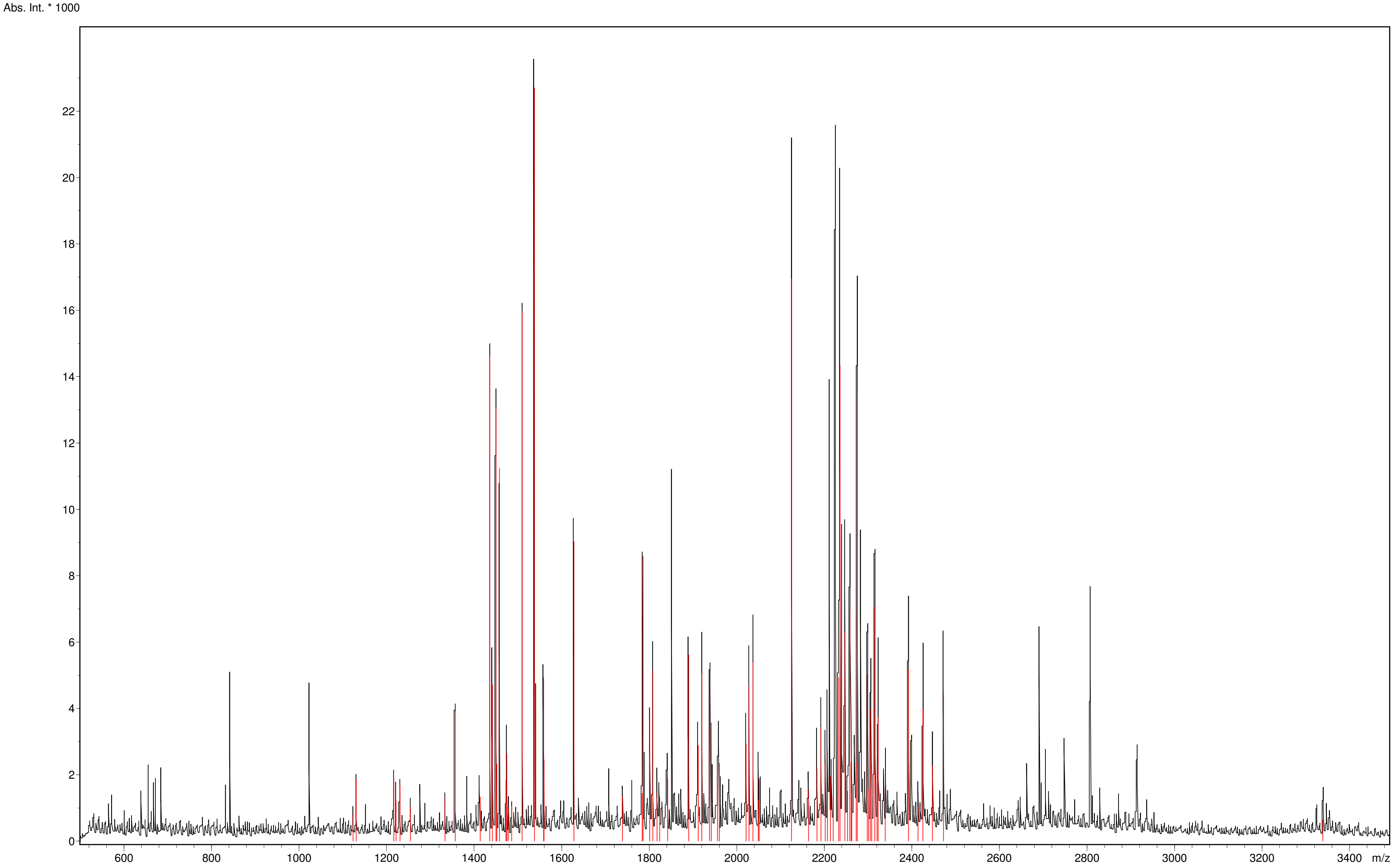


 Copyright © 1999-2008 Bruker Daltonik GmbH 

#### Spectrum Analysis Report
### Rieske subunit

**Sequence data:**

Intensity Coverage: 1.1 % (3580 cnts)
Sequence Coverage MS: 13.0%
Sequence Coverage MS/MS: 0.0%
pI (isoelectric point): 9.3

| 10 | 20 | 30 | 40 | 50 | 60 | 70 | 80 | 90 | 100 | 110 | 120 |
|---|---|---|---|---|---|---|---|---|---|---|---|
| MLGIRSSVKT | CFKPMSLTSK | RLISQSLLAS | KSTYRTPNFD | DVLKENNDAD | KGRSYAYFMV | GAMGLLSSAG | AKSTVETFIS | SMTATADVLA | MAKVEVNLAA | IPLGKNVVVK | WQGKPVFIRH |
| 130 | 140 | 150 | 160 | 170 | 180 | 190 | 200 | 210 | 220 | | |
| RTPHEIQEAN | SVDMSALKDP | QTDADRVKDP | QWLIMLGICT | HLGCVPIGEA | GDFGGWFCPC | HGSHYDISGR | IRKGPAPLNL | EIPAYEFDGD | KVIVG | | |

**Display Parameter:**

MH+ (mono): 1.008 MH+ (avg): 1.008 Threshold (a.i.): 0.000 Tolerance (Da): 0.500
Number of Peaks: 74

**Peaklist:**

| Peak | Mass | Intensity | Peak | Mass | Intensity | Peak | Mass | Intensity | Peak | Mass | Intensity | Peak | Mass | Intensity |
|---|---|---|---|---|---|---|---|---|---|---|---|---|---|---|
| 1 | 1123.583 | 808.568 | 2 | 1130.622 | 1869.448 | 3 | 1216.596 | 1883.277 | 4 | 1221.606 | 1226.315 | 5 | 1230.633 | 1677.819 |
| 6 | 1254.531 | 1032.225 | 7 | 1333.672 | 1337.183 | 8 | 1356.622 | 3905.019 | 9 | 1413.702 | 1328.696 | 10 | 1435.724 | 14621.908 |
| 11 | 1441.744 | 4709.528 | 12 | 1449.741 | 13033.796 | 13 | 1451.725 | 2323.019 | 14 | 1457.727 | 11235.150 | 15 | 1472.716 | 1826.124 |
| 16 | 1474.723 | 2623.582 | 17 | 1477.723 | 971.750 | 18 | 1485.749 | 1046.601 | 19 | 1509.671 | 15966.638 | 20 | 1537.725 | 22703.080 |
| 21 | 1558.823 | 4932.817 | 22 | 1559.714 | 2454.025 | 23 | 1627.815 | 9030.708 | 24 | 1738.836 | 1349.505 | 25 | 1783.819 | 1440.397 |
| 26 | 1785.965 | 8593.898 | 27 | 1800.915 | 3643.763 | 28 | 1807.949 | 5120.049 | 29 | 1817.848 | 1461.242 | 30 | 1823.906 | 1323.379 |
| 31 | 1841.902 | 1891.399 | 32 | 1890.059 | 5611.720 | 33 | 1912.043 | 2878.011 | 34 | 1919.942 | 5041.679 | 35 | 1937.979 | 4709.868 |
| 36 | 1941.934 | 3570.011 | 37 | 1955.951 | 1710.427 | 38 | 1959.966 | 2346.213 | 39 | 2021.032 | 2934.943 | 40 | 2027.950 | 4711.074 |
| 41 | 2037.024 | 5380.085 | 42 | 2048.923 | 1236.841 | 43 | 2051.005 | 1878.804 | 44 | 2125.105 | 16947.955 | 45 | 2164.078 | 1550.771 |
| 46 | 2183.020 | 2179.902 | 47 | 2192.118 | 3430.623 | 48 | 2202.102 | 2358.338 | 49 | 2207.097 | 2693.954 | 50 | 2214.134 | 1944.392 |
| 51 | 2220.090 | 1328.941 | 52 | 2233.079 | 4933.313 | 53 | 2236.103 | 14340.864 | 54 | 2247.118 | 6304.303 | 55 | 2253.096 | 1507.950 |
| 56 | 2258.092 | 6961.212 | 57 | 2261.169 | 2382.070 | 58 | 2262.661 | 1441.921 | 59 | 2273.062 | 2367.695 | 60 | 2275.191 | 13389.789 |
| 61 | 2298.170 | 5004.297 | 62 | 2301.153 | 1564.191 | 63 | 2305.138 | 3968.486 | 64 | 2308.101 | 2283.139 | 65 | 2315.167 | 7059.208 |
| 66 | 2320.158 | 2085.198 | 67 | 2323.096 | 3727.940 | 68 | 2339.098 | 1628.892 | 69 | 2392.209 | 5182.606 | 70 | 2414.193 | 1030.947 |
| 71 | 2425.200 | 4004.982 | 72 | 2447.172 | 2280.361 | 73 | 2472.270 | 4390.654 | 74 | 3338.700 | 645.613 | | | |


Copyright © 1999-2008 Bruker Daltonik GmbH

#### Spectrum Analysis Report
### iso-1 Cc

| | | | | | | | |
|---|---|---|---|---|---|---|---|
| Sequence Name: | | Formula: | | Parentmass: | | Mass Error: | |
| MH+ (mono): | 1.008 | MH+ (avg): | 1.008 | Threshold (a.i.): | 0.000 | Tolerance (Da): | 0.500 |
| Number of Peaks: | 74 | Above Threshold: | | Assigned Peaks: | | Not assigned Peaks: | |

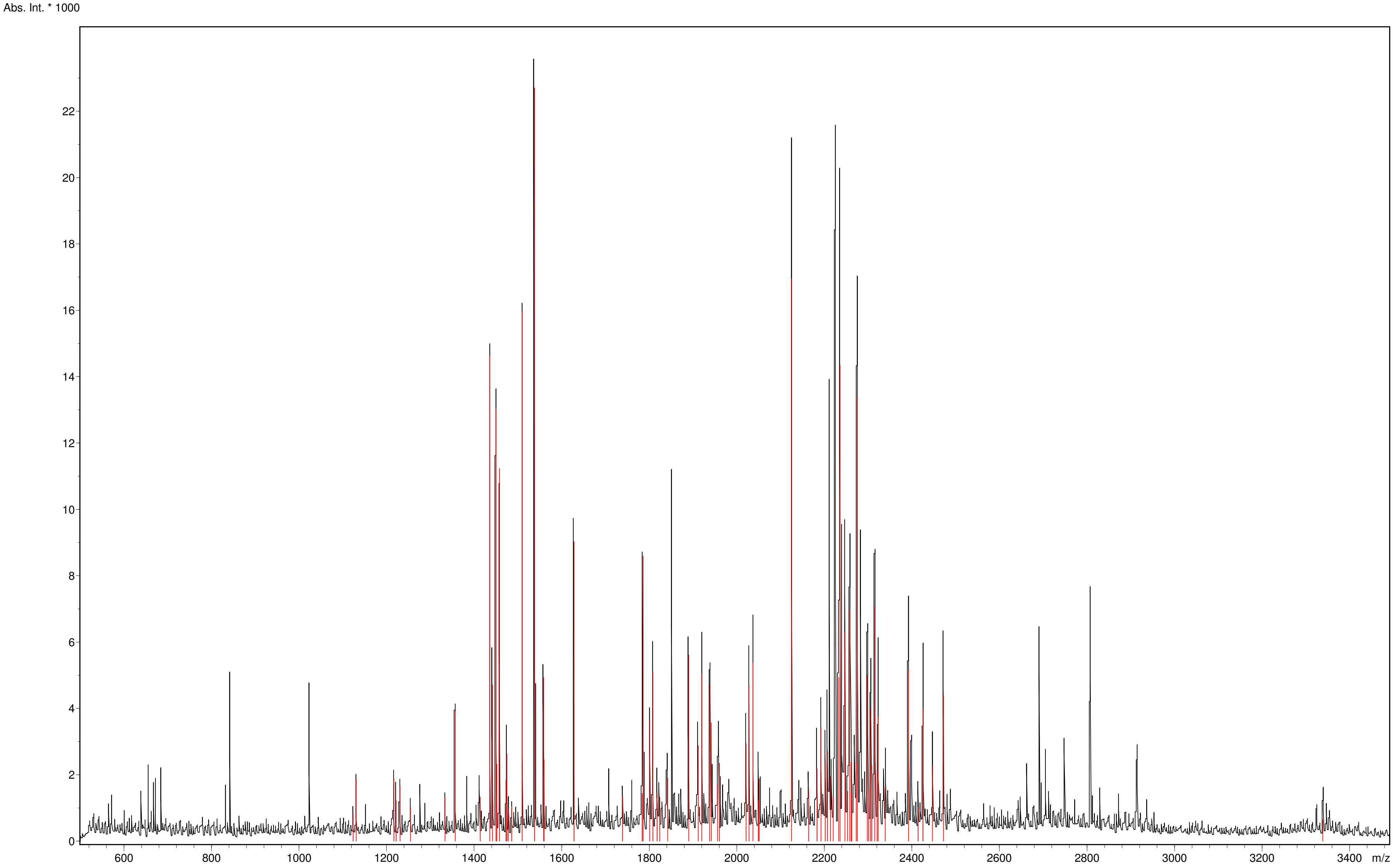



Copyright © 1999-2008 Bruker Daltonik GmbH

### Spectrum Analysis Report
#### iso-1 Cc

**Sequence data:**

| | | | |
|---|---|---|---|
| Intensity Coverage: | 1.7 % (5357 cnts) | Sequence Coverage MS: | 17.4% |
| Sequence Coverage MS/MS: | 0.0% | pI (isoelectric point): | 10.1 |

| 10 | 20 | 30 | 40 | 50 | 60 | 70 | 80 | 90 | 100 | 110 |
|---|---|---|---|---|---|---|---|---|---|---|
| MTEFKAGSAK | KGATLFKTRC | LQCHTVEKGG | PHKVGPNLHG | IFGRHSGQAE | GYSYTDANIK | KNVLWDENNM | SEYLTNPKKY | IPGTKMAFGG | LKKEKDRNDL | ITYLKKACE |

**Display Parameter:**

| | | | | | | | |
|---|---|---|---|---|---|---|---|
| MH+ (mono): | 1.008 | MH+ (avg): | 1.008 | Threshold (a.i.): | 0.000 | Tolerance (Da): | 0.500 |
| Number of Peaks: | 74 | | | | | | |

**Peaklist:**

| Peak | Mass | Intensity | Peak | Mass | Intensity | Peak | Mass | Intensity | Peak | Mass | Intensity | Peak | Mass | Intensity |
|---|---|---|---|---|---|---|---|---|---|---|---|---|---|---|
| 1 | 1123.583 | 808.568 | 2 | 1130.622 | 1869.448 | 3 | 1216.596 | 1883.277 | 4 | 1221.606 | 1226.315 | 5 | 1230.633 | 1677.819 |
| 6 | 1254.531 | 1032.225 | 7 | 1333.672 | 1337.183 | 8 | 1356.622 | 3905.019 | 9 | 1413.702 | 1328.696 | 10 | 1435.724 | 14621.908 |
| 11 | 1441.744 | 4709.528 | 12 | 1449.741 | 13033.796 | 13 | 1451.725 | 2323.019 | 14 | 1457.727 | 11235.150 | 15 | 1472.716 | 1826.124 |
| 16 | 1474.723 | 2623.582 | 17 | 1477.723 | 971.750 | 18 | 1485.749 | 1046.601 | 19 | 1509.671 | 15966.638 | 20 | 1537.725 | 22703.080 |
| 21 | 1558.823 | 4932.817 | 22 | 1559.714 | 2454.025 | 23 | 1627.815 | 9030.708 | 24 | 1738.836 | 1349.505 | 25 | 1783.819 | 1440.397 |
| 26 | 1785.965 | 8593.898 | 27 | 1800.915 | 3643.763 | 28 | 1807.949 | 5120.049 | 29 | 1817.848 | 1461.242 | 30 | 1823.906 | 1323.379 |
| 31 | 1841.902 | 1891.399 | 32 | 1890.059 | 5611.720 | 33 | 1912.043 | 2878.011 | 34 | 1919.942 | 5041.679 | 35 | 1937.979 | 4709.868 |
| 36 | 1941.934 | 3570.011 | 37 | 1955.951 | 1710.427 | 38 | 1959.966 | 2346.213 | 39 | 2021.032 | 2934.943 | 40 | 2027.950 | 4711.074 |
| 41 | 2037.024 | 5380.085 | 42 | 2048.923 | 1236.841 | 43 | 2051.005 | 1878.804 | 44 | 2125.105 | 16947.955 | 45 | 2164.078 | 1550.771 |
| 46 | 2183.020 | 2179.902 | 47 | 2192.118 | 3430.623 | 48 | 2202.102 | 2358.338 | 49 | 2207.097 | 2693.954 | 50 | 2214.134 | 1944.392 |
| 51 | 2220.090 | 1328.941 | 52 | 2233.079 | 4933.313 | 53 | 2236.103 | 14340.864 | 54 | 2247.118 | 6304.303 | 55 | 2253.096 | 1507.950 |
| 56 | 2258.092 | 6961.212 | 57 | 2261.169 | 2382.070 | 58 | 2262.661 | 1441.921 | 59 | 2273.062 | 2367.695 | 60 | 2275.191 | 13389.789 |
| 61 | 2298.170 | 5004.297 | 62 | 2301.153 | 1564.191 | 63 | 2305.138 | 3968.486 | 64 | 2308.101 | 2283.139 | 65 | 2315.167 | 7059.208 |
| 66 | 2320.158 | 2085.198 | 67 | 2323.096 | 3727.940 | 68 | 2339.098 | 1628.892 | 69 | 2392.209 | 5182.606 | 70 | 2414.193 | 1030.947 |
| 71 | 2425.200 | 4004.982 | 72 | 2447.172 | 2280.361 | 73 | 2472.270 | 4390.654 | 74 | 3338.700 | 645.613 | | | |


Copyright © 1999-2008 Bruker Daltonik GmbH

#### Spectrum Analysis Report
#### iso-2 Cc

| | | | | | | | |
|---|---|---|---|---|---|---|---|
| Sequence Name: | | Formula: | | Parentmass: | | Mass Error: | |
| MH+ (mono): | 1.008 | MH+ (avg): | 1.008 | Threshold (a.i.): | 0.000 | Tolerance (Da): | 0.500 |
| Number of Peaks: | 74 | Above Threshold: | | Assigned Peaks: | | Not assigned Peaks: | |

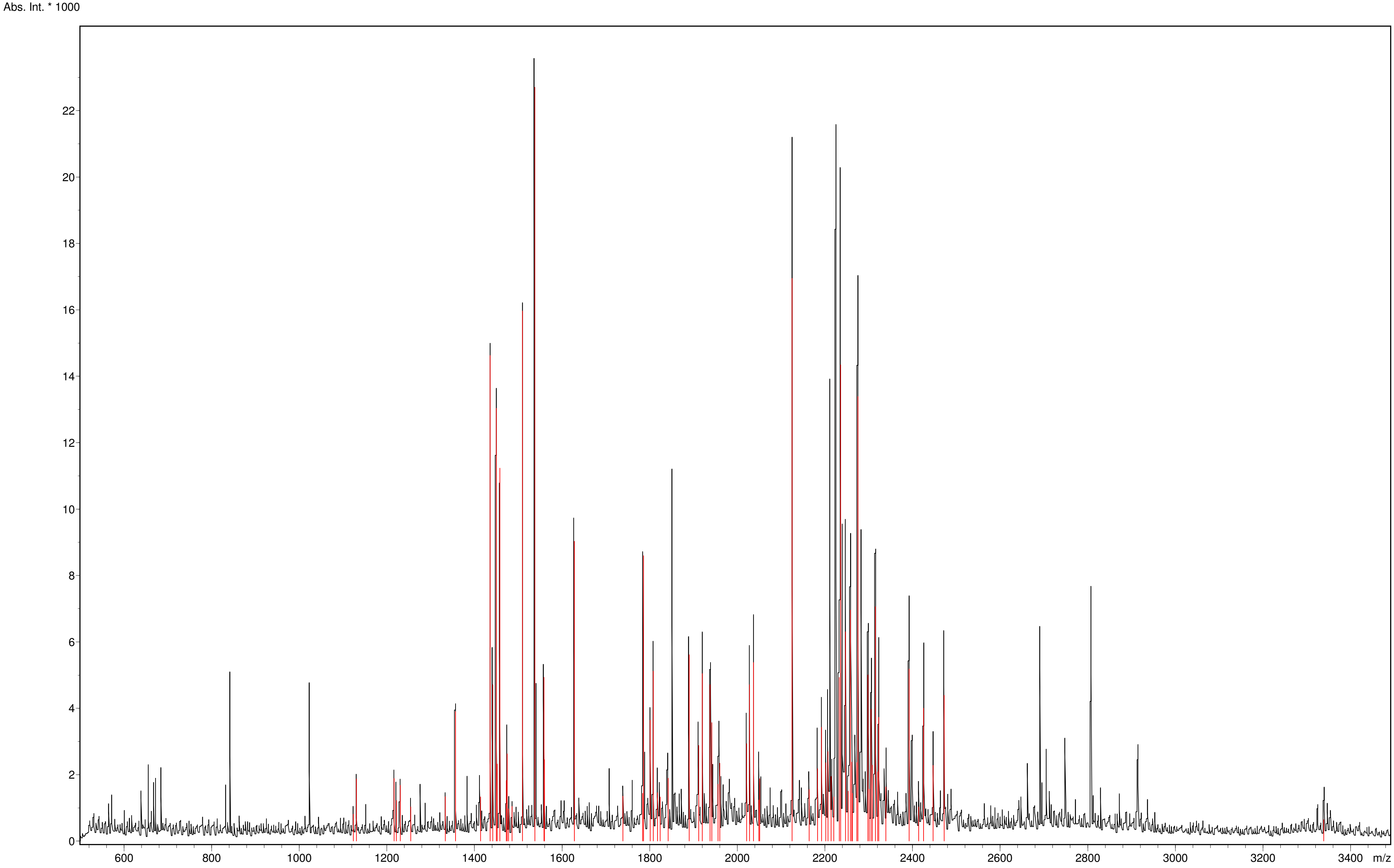



Copyright © 1999-2008 Bruker Daltonik GmbH

#### Spectrum Analysis Report
### iso-2 Cc

**Sequence data:**

| | | | |
|---|---|---|---|
| Intensity Coverage: | 1.1 % (0 cnts) | Sequence Coverage MS: | 0.0% |
| Sequence Coverage MS/MS: | 0.0% | pI (isoelectric point): | 10.2 |

| 10 | 20 | 30 | 40 | 50 | 60 | 70 | 80 | 90 | 100 | 110 | 120 |
|---|---|---|---|---|---|---|---|---|---|---|---|
| MAKESTGFKP | GSAKKGATLF | KTRCQQCHTI | EEGGPNKVGP | NLHGIFGRHS | GQVKGYSYTD | ANINKNVKWD | EDSMSEYLTN | PKKYIPGTKM | AFAGLKKEKD | RNDLITYMTK | AAK |

**Display Parameter:**

| | | | | | | | |
|---|---|---|---|---|---|---|---|
| MH+ (mono): | 1.008 | MH+ (avg): | 1.008 | Threshold (a.i.): | 0.000 | Tolerance (Da): | 0.500 |
| Number of Peaks: | 74 | | | | | | |

**Peaklist:**

| Peak | Mass | Intensity | Peak | Mass | Intensity | Peak | Mass | Intensity | Peak | Mass | Intensity | Peak | Mass | Intensity |
|---|---|---|---|---|---|---|---|---|---|---|---|---|---|---|
| 1 | 1123.583 | 808.568 | 2 | 1130.622 | 1869.448 | 3 | 1216.596 | 1883.277 | 4 | 1221.606 | 1226.315 | 5 | 1230.633 | 1677.819 |
| 6 | 1254.531 | 1032.225 | 7 | 1333.672 | 1337.183 | 8 | 1356.622 | 3905.019 | 9 | 1413.702 | 1328.696 | 10 | 1435.724 | 14621.908 |
| 11 | 1441.744 | 4709.528 | 12 | 1449.741 | 13033.796 | 13 | 1451.725 | 2323.019 | 14 | 1457.727 | 11235.150 | 15 | 1472.716 | 1826.124 |
| 16 | 1474.723 | 2623.582 | 17 | 1477.723 | 971.750 | 18 | 1485.749 | 1046.601 | 19 | 1509.671 | 15966.638 | 20 | 1537.725 | 22703.080 |
| 21 | 1558.823 | 4932.817 | 22 | 1559.714 | 2454.025 | 23 | 1627.815 | 9030.708 | 24 | 1738.836 | 1349.505 | 25 | 1783.819 | 1440.397 |
| 26 | 1785.965 | 8593.898 | 27 | 1800.915 | 3643.763 | 28 | 1807.949 | 5120.049 | 29 | 1817.848 | 1461.242 | 30 | 1823.906 | 1323.379 |
| 31 | 1841.902 | 1891.399 | 32 | 1890.059 | 5611.720 | 33 | 1912.043 | 2878.011 | 34 | 1919.942 | 5041.679 | 35 | 1937.979 | 4709.868 |
| 36 | 1941.934 | 3570.011 | 37 | 1955.951 | 1710.427 | 38 | 1959.966 | 2346.213 | 39 | 2021.032 | 2934.943 | 40 | 2027.950 | 4711.074 |
| 41 | 2037.024 | 5380.085 | 42 | 2048.923 | 1236.841 | 43 | 2051.005 | 1878.804 | 44 | 2125.105 | 16947.955 | 45 | 2164.078 | 1550.771 |
| 46 | 2183.020 | 2179.902 | 47 | 2192.118 | 3430.623 | 48 | 2202.102 | 2358.338 | 49 | 2207.097 | 2693.954 | 50 | 2214.134 | 1944.392 |
| 51 | 2220.090 | 1328.941 | 52 | 2233.079 | 4933.313 | 53 | 2236.103 | 14340.864 | 54 | 2247.118 | 6304.303 | 55 | 2253.096 | 1507.950 |
| 56 | 2258.092 | 6961.212 | 57 | 2261.169 | 2382.070 | 58 | 2262.661 | 1441.921 | 59 | 2273.062 | 2367.695 | 60 | 2275.191 | 13389.789 |
| 61 | 2298.170 | 5004.297 | 62 | 2301.153 | 1564.191 | 63 | 2305.138 | 3968.486 | 64 | 2308.101 | 2283.139 | 65 | 2315.167 | 7059.208 |
| 66 | 2320.158 | 2085.198 | 67 | 2323.096 | 3727.940 | 68 | 2339.098 | 1628.892 | 69 | 2392.209 | 5182.606 | 70 | 2414.193 | 1030.947 |
| 71 | 2425.200 | 4004.982 | 72 | 2447.172 | 2280.361 | 73 | 2472.270 | 4390.654 | 74 | 3338.700 | 645.613 | | | |


Copyright © 1999-2008 Bruker Daltonik GmbH

**Unveiling the role of yeast cytochrome *c* isoforms in the assembly of mitochondrial supercomplexes and the control of respiratory chain rate**

#### ANNEX C. TRYPTIC DIGESTION ANALYSES OF BN-PAGE

### YPD-2

#### Spectrum Analysis Report

### Cytochrome c1

| | | | | | | | |
|---|---|---|---|---|---|---|---|
| Sequence Name: | | Formula: | | Parentmass: | | Mass Error: | |
| MH+ (mono): | 1.008 | MH+ (avg): | 1.008 | Threshold (a.i.): | 0.000 | Tolerance (Da): | 0.500 |
| Number of Peaks: | 51 | Above Threshold: | | Assigned Peaks: | | Not assigned Peaks: | |

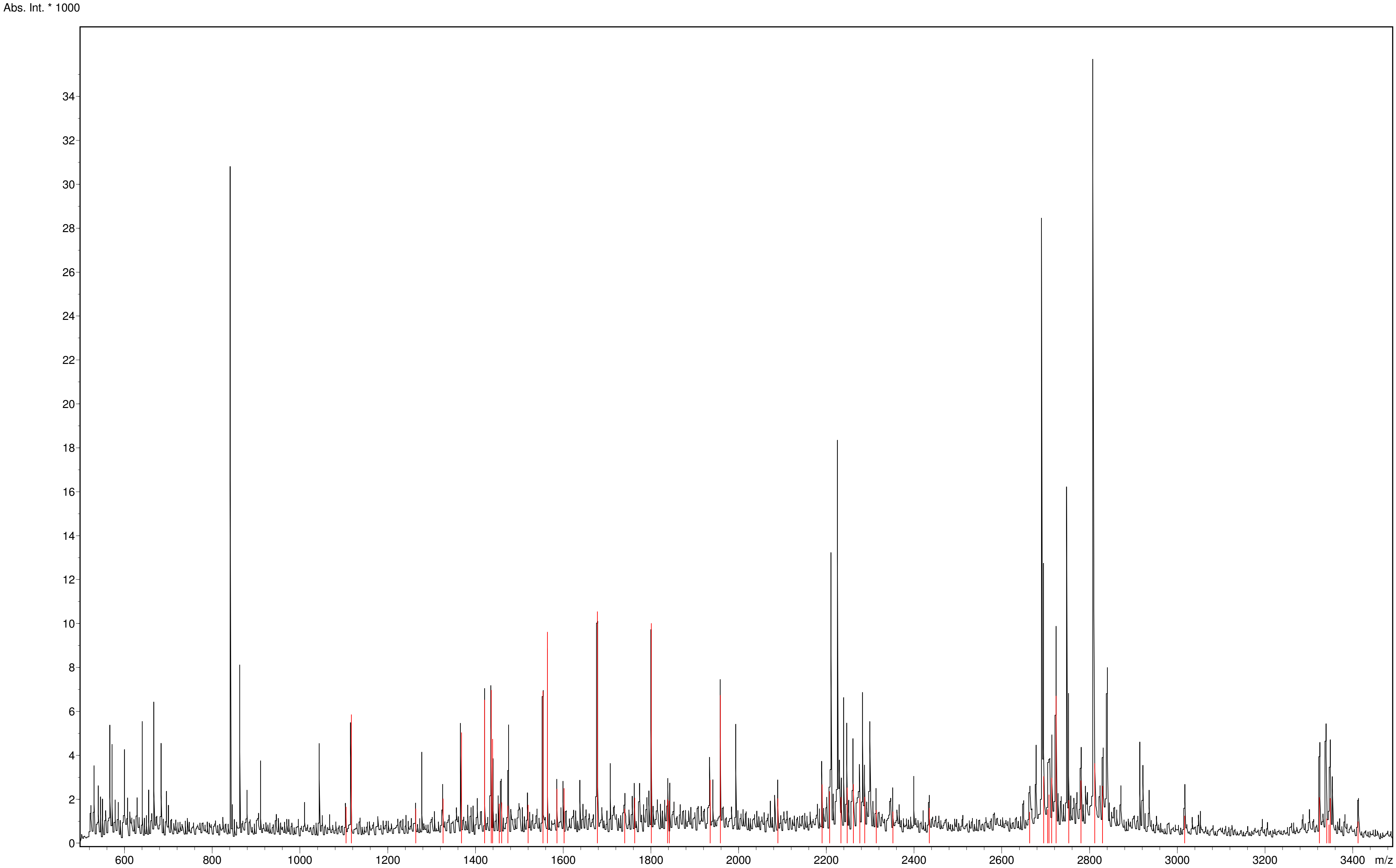



Copyright © 1999-2008 Bruker Daltonik GmbH

### Spectrum Analysis Report
#### Cytochrome c1

**Sequence data:**

| | | | |
|---|---|---|---|
| Intensity Coverage: | 1.6 % (2520 cnts) | Sequence Coverage MS: | 5.8% |
| Sequence Coverage MS/MS: | 0.0% | pI (isoelectric point): | 9.0 |

| 10 | 20 | 30 | 40 | 50 | 60 | 70 | 80 | 90 | 100 | 110 | 120 |
|---|---|---|---|---|---|---|---|---|---|---|---|
| MFSNLSKRWA | QRTLSKSFYS | TATGAASKSG | KLTQKLVTAG | VAAAGITAST | LLYADSLTAE | AMTAAEHGLH | APAYAWSHNG | PFETFDHASI | RRGYQVYREV | CAACHSLDRV | AWRTLVGVSH |
| **130** | **140** | **150** | **160** | **170** | **180** | **190** | **200** | **210** | **220** | **230** | **240** |
| TNEEVRNMAE | EFEYDDEPDE | QGNPKKRPGK | LSDYIPGPYP | NEQAARAANQ | GALPPDLSLI | VKARHGGCDY | IFSLLTGYPD | EPPAGVALPP | GSNYNPYFPG | GSIAMARVLF | DDMVEYEDGT |
| **250** | **260** | **270** | **280** | **290** | **300** | **310** | | | | | |
| PATTSQMAKD | VTTFLNWCAE | PEHDERKRLG | LKTVIILSSL | YLLSIWVKKF | KWAGIKTRKF | VFNPPKPRK | | | | | |

**Display Parameter:**

| | | | | | | | |
|---|---|---|---|---|---|---|---|
| MH+ (mono): | 1.008 | MH+ (avg): | 1.008 | Threshold (a.i.): | 0.000 | Tolerance (Da): | 0.500 |
| Number of Peaks: | 51 | | | | | | |

**Peaklist:**

| Peak | Mass | Intensity | Peak | Mass | Intensity | Peak | Mass | Intensity | Peak | Mass | Intensity | Peak | Mass | Intensity |
|---|---|---|---|---|---|---|---|---|---|---|---|---|---|---|
| 1 | 1104.560 | 1642.571 | 2 | 1116.630 | 5849.584 | 3 | 1263.663 | 1558.691 | 4 | 1325.666 | 2021.085 | 5 | 1367.713 | 5031.395 |
| 6 | 1420.660 | 6520.024 | 7 | 1435.711 | 6964.991 | 8 | 1438.808 | 4732.983 | 9 | 1453.751 | 1787.974 | 10 | 1459.697 | 1839.918 |
| 11 | 1473.744 | 1692.960 | 12 | 1519.716 | 1731.683 | 13 | 1553.689 | 6907.891 | 14 | 1563.820 | 9604.605 | 15 | 1585.765 | 2458.610 |
| 16 | 1601.761 | 2493.508 | 17 | 1677.868 | 10543.567 | 18 | 1739.825 | 1547.629 | 19 | 1762.932 | 2058.760 | 20 | 1800.868 | 10007.568 |
| 21 | 1837.860 | 1991.144 | 22 | 1841.868 | 1923.329 | 23 | 1934.999 | 2868.493 | 24 | 1958.062 | 6724.025 | 25 | 2089.050 | 2031.296 |
| 26 | 2190.081 | 2662.029 | 27 | 2207.120 | 2349.230 | 28 | 2233.033 | 2087.487 | 29 | 2247.084 | 2520.425 | 30 | 2261.145 | 2671.969 |
| 31 | 2276.148 | 2018.556 | 32 | 2287.098 | 2098.454 | 33 | 2313.207 | 1504.250 | 34 | 2351.162 | 1426.172 | 35 | 2434.370 | 1590.964 |
| 36 | 2663.371 | 1413.873 | 37 | 2695.326 | 3034.053 | 38 | 2704.263 | 1533.906 | 39 | 2707.327 | 2197.558 | 40 | 2713.300 | 2969.736 |
| 41 | 2723.317 | 6695.497 | 42 | 2752.357 | 1946.806 | 43 | 2780.337 | 2842.090 | 44 | 2811.361 | 3619.950 | 45 | 2829.327 | 2595.014 |
| 46 | 3016.614 | 1242.117 | 47 | 3323.696 | 2077.063 | 48 | 3340.706 | 1103.996 | 49 | 3345.629 | 830.477 | 50 | 3348.564 | 2038.476 |
| 51 | 3411.646 | 975.535 | | | | | | | | | | | | |


Copyright © 1999-2008 Bruker Daltonik GmbH

#### Spectrum Analysis Report
### Rieske subunit

| | | | | | | | |
|---|---|---|---|---|---|---|---|
| Sequence Name: | | Formula: | | Parentmass: | | Mass Error: | |
| MH+ (mono): | 1.008 | MH+ (avg): | 1.008 | Threshold (a.i.): | 0.000 | Tolerance (Da): | 0.500 |
| Number of Peaks: | 51 | Above Threshold: | | Assigned Peaks: | | Not assigned Peaks: | |

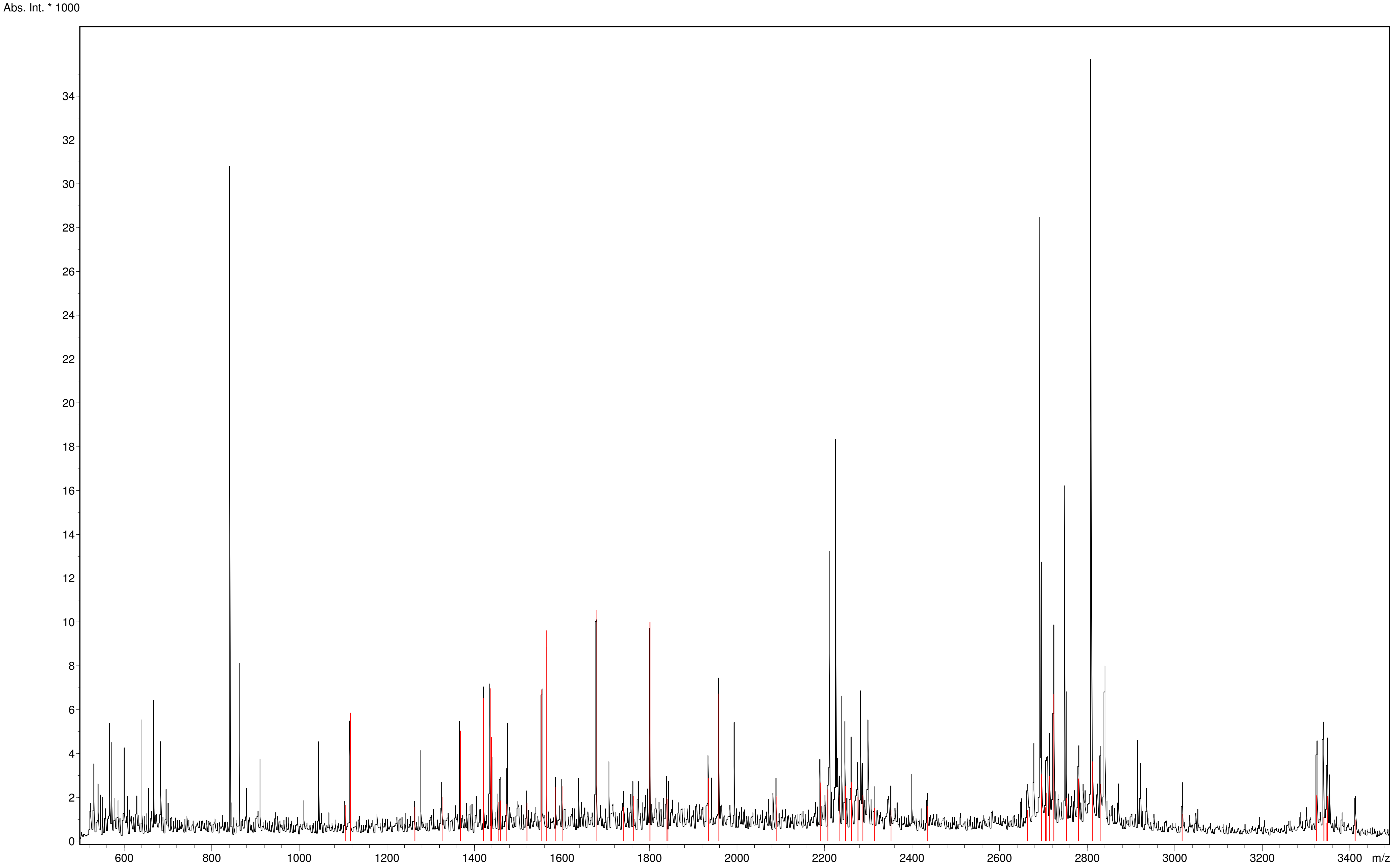



Copyright © 1999-2008 Bruker Daltonik GmbH

#### Spectrum Analysis Report
### Rieske subunit

**Sequence data:**

| | | | |
|---|---|---|---|
| Intensity Coverage: | 3.0 % (4760 cnts) | Sequence Coverage MS: | 19.1% |
| Sequence Coverage MS/MS: | 0.0% | pI (isoelectric point): | 9.3 |

| 10 | 20 | 30 | 40 | 50 | 60 | 70 | 80 | 90 | 100 | 110 | 120 |
|---|---|---|---|---|---|---|---|---|---|---|---|
| MLGIRSSVKT | CFKPMSLTSK | RLISQSLLAS | KSTYRTPNFD | DVLKENNDAD | KGRSYAYFMV | GAMGLLSSAG | AKSTVETFIS | SMTATADVLA | MAKVEVNLAA | IPLGKNVVVK | WQGKPVFIRH |
| **130** | **140** | **150** | **160** | **170** | **180** | **190** | **200** | **210** | **220** | | |
| RTPHEIQEAN | SVDMSALKDP | QTDADRVKDP | QWLIMLGICT | HLGCVPIGEA | GDFGGWFCPC | HGSHYDISGR | IRKGPAPLNL | EIPAYEFDGD | KVIVG | | |

**Display Parameter:**

| | | | | | | | |
|---|---|---|---|---|---|---|---|
| MH+ (mono): | 1.008 | MH+ (avg): | 1.008 | Threshold (a.i.): | 0.000 | Tolerance (Da): | 0.500 |
| Number of Peaks: | 51 | | | | | | |

**Peaklist:**

| Peak | Mass | Intensity | Peak | Mass | Intensity | Peak | Mass | Intensity | Peak | Mass | Intensity | Peak | Mass | Intensity |
|---|---|---|---|---|---|---|---|---|---|---|---|---|---|---|
| 1 | 1104.560 | 1642.571 | 2 | 1116.630 | 5849.584 | 3 | 1263.663 | 1558.691 | 4 | 1325.666 | 2021.085 | 5 | 1367.713 | 5031.395 |
| 6 | 1420.660 | 6520.024 | 7 | 1435.711 | 6964.991 | 8 | 1438.808 | 4732.983 | 9 | 1453.751 | 1787.974 | 10 | 1459.697 | 1839.918 |
| 11 | 1473.744 | 1692.960 | 12 | 1519.716 | 1731.683 | 13 | 1553.689 | 6907.891 | 14 | 1563.820 | 9604.605 | 15 | 1585.765 | 2458.610 |
| 16 | 1601.761 | 2493.508 | 17 | 1677.868 | 10543.567 | 18 | 1739.825 | 1547.629 | 19 | 1762.932 | 2058.760 | 20 | 1800.868 | 10007.568 |
| 21 | 1837.860 | 1991.144 | 22 | 1841.868 | 1923.329 | 23 | 1934.999 | 2868.493 | 24 | 1958.062 | 6724.025 | 25 | 2089.050 | 2031.296 |
| 26 | 2190.081 | 2662.029 | 27 | 2207.120 | 2349.230 | 28 | 2233.033 | 2087.487 | 29 | 2247.084 | 2520.425 | 30 | 2261.145 | 2671.969 |
| 31 | 2276.148 | 2018.556 | 32 | 2287.098 | 2098.454 | 33 | 2313.207 | 1504.250 | 34 | 2351.162 | 1426.172 | 35 | 2434.370 | 1590.964 |
| 36 | 2663.371 | 1413.873 | 37 | 2695.326 | 3034.053 | 38 | 2704.263 | 1533.906 | 39 | 2707.327 | 2197.558 | 40 | 2713.300 | 2969.736 |
| 41 | 2723.317 | 6695.497 | 42 | 2752.357 | 1946.806 | 43 | 2780.337 | 2842.090 | 44 | 2811.361 | 3619.950 | 45 | 2829.327 | 2595.014 |
| 46 | 3016.614 | 1242.117 | 47 | 3323.696 | 2077.063 | 48 | 3340.706 | 1103.996 | 49 | 3345.629 | 830.477 | 50 | 3348.564 | 2038.476 |
| 51 | 3411.646 | 975.535 | | | | | | | | | | | | |


Copyright © 1999-2008 Bruker Daltonik GmbH

**Spectrum Analysis Report**

#### iso-1 Cc

| | | | | | | | |
|---|---|---|---|---|---|---|---|
| Sequence Name: | | Formula: | | Parentmass: | | Mass Error: | |
| MH+ (mono): | 1.008 | MH+ (avg): | 1.008 | Threshold (a.i.): | 0.000 | Tolerance (Da): | 0.500 |
| Number of Peaks: | 51 | Above Threshold: | | Assigned Peaks: | | Not assigned Peaks: | |

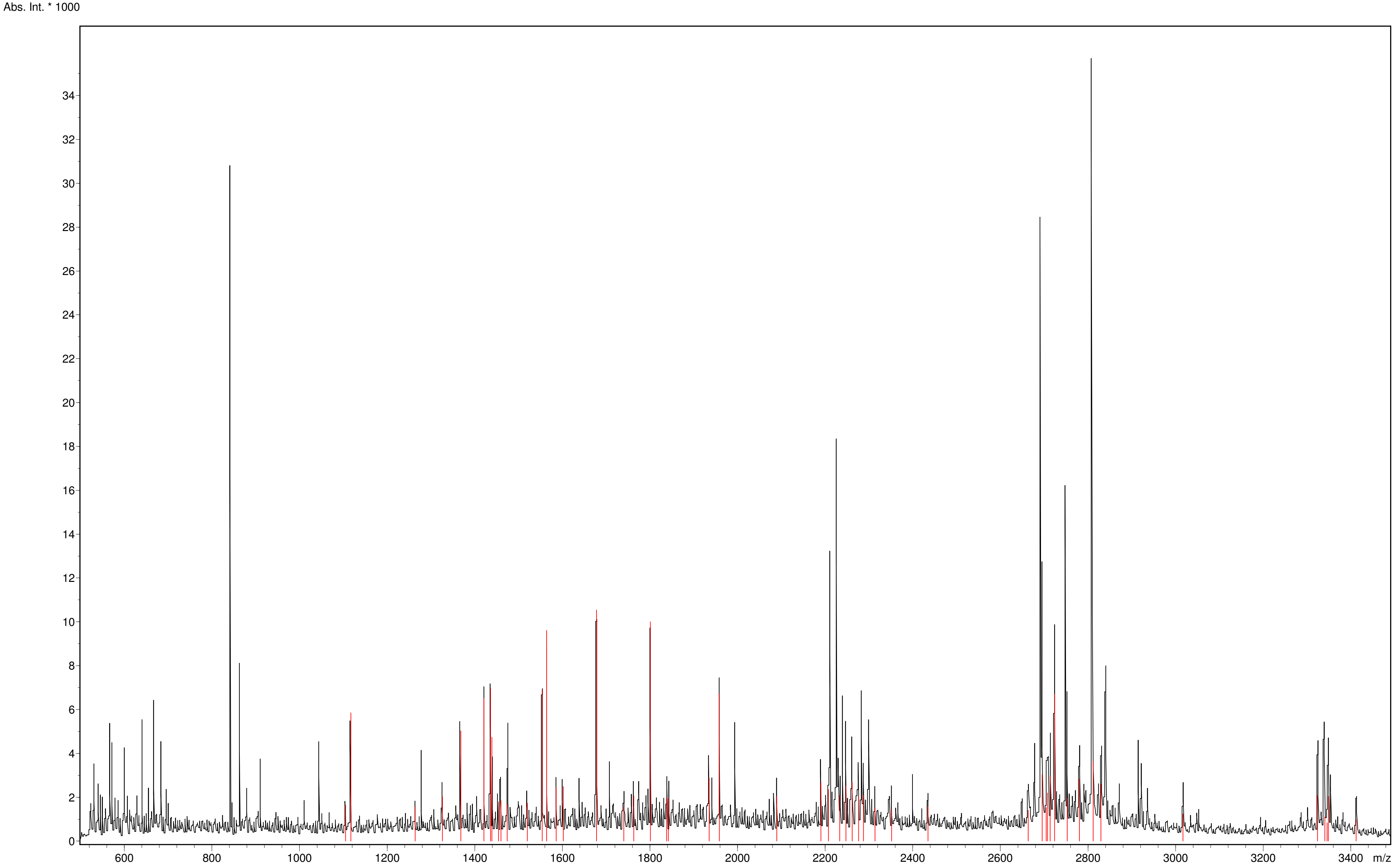



Copyright © 1999-2008 Bruker Daltonik GmbH

#### Spectrum Analysis Report

### iso-1 Cc

**Sequence data:**

| | | | |
|---|---|---|---|
| Intensity Coverage: | 0.8 % (1242 cnts) | Sequence Coverage MS: | 25.7% |
| Sequence Coverage MS/MS: | 0.0% | pI (isoelectric point): | 10.1 |

```
        10         20         30         40         50         60         70         80         90        100        110
MTEFKAGSAK KGATLFKTRC LQCHTVEKGG PHKVGPNLHG IFGRHSGQAE GYSYTDANIK KNVLWDENNM SEYLTNPKKY IPGTKMAFGG LKKEKDRNDL ITYLKKACE
```

**Display Parameter:**

| | | | | | | | |
|---|---|---|---|---|---|---|---|
| MH+ (mono): | 1.008 | MH+ (avg): | 1.008 | Threshold (a.i.): | 0.000 | Tolerance (Da): | 0.500 |
| Number of Peaks: | 51 | | | | | | |

**Peaklist:**

| Peak | Mass | Intensity | Peak | Mass | Intensity | Peak | Mass | Intensity | Peak | Mass | Intensity | Peak | Mass | Intensity |
|---|---|---|---|---|---|---|---|---|---|---|---|---|---|---|
| 1 | 1104.560 | 1642.571 | 2 | 1116.630 | 5849.584 | 3 | 1263.663 | 1558.691 | 4 | 1325.666 | 2021.085 | 5 | 1367.713 | 5031.395 |
| 6 | 1420.660 | 6520.024 | 7 | 1435.711 | 6964.991 | 8 | 1438.808 | 4732.983 | 9 | 1453.751 | 1787.974 | 10 | 1459.697 | 1839.918 |
| 11 | 1473.744 | 1692.960 | 12 | 1519.716 | 1731.683 | 13 | 1553.689 | 6907.891 | 14 | 1563.820 | 9604.605 | 15 | 1585.765 | 2458.610 |
| 16 | 1601.761 | 2493.508 | 17 | 1677.868 | 10543.567 | 18 | 1739.825 | 1547.629 | 19 | 1762.932 | 2058.760 | 20 | 1800.868 | 10007.568 |
| 21 | 1837.860 | 1991.144 | 22 | 1841.868 | 1923.329 | 23 | 1934.999 | 2868.493 | 24 | 1958.062 | 6724.025 | 25 | 2089.050 | 2031.296 |
| 26 | 2190.081 | 2662.029 | 27 | 2207.120 | 2349.230 | 28 | 2233.033 | 2087.487 | 29 | 2247.084 | 2520.425 | 30 | 2261.145 | 2671.969 |
| 31 | 2276.148 | 2018.556 | 32 | 2287.098 | 2098.454 | 33 | 2313.207 | 1504.250 | 34 | 2351.162 | 1426.172 | 35 | 2434.370 | 1590.964 |
| 36 | 2663.371 | 1413.873 | 37 | 2695.326 | 3034.053 | 38 | 2704.263 | 1533.906 | 39 | 2707.327 | 2197.558 | 40 | 2713.300 | 2969.736 |
| 41 | 2723.317 | 6695.497 | 42 | 2752.357 | 1946.806 | 43 | 2780.337 | 2842.090 | 44 | 2811.361 | 3619.950 | 45 | 2829.327 | 2595.014 |
| 46 | 3016.614 | 1242.117 | 47 | 3323.696 | 2077.063 | 48 | 3340.706 | 1103.996 | 49 | 3345.629 | 830.477 | 50 | 3348.564 | 2038.476 |
| 51 | 3411.646 | 975.535 | | | | | | | | | | | | |


Copyright © 1999-2008 Bruker Daltonik GmbH

**Unveiling the role of yeast cytochrome *c* isoforms in the assembly of mitochondrial supercomplexes and the control of respiratory chain rate**

#### ANNEX C. TRYPTIC DIGESTION ANALYSES OF BN-PAGE

### YPG-1

#### Spectrum Analysis Report
### Cytochrome c1

| | | | | | | | |
|---|---|---|---|---|---|---|---|
| Sequence Name: | | Formula: | | Parentmass: | | Mass Error: | |
| MH+ (mono): | 1.008 | MH+ (avg): | 1.008 | Threshold (a.i.): | 0.000 | Tolerance (Da): | 0.500 |
| Number of Peaks: | 59 | Above Threshold: | | Assigned Peaks: | | Not assigned Peaks: | |

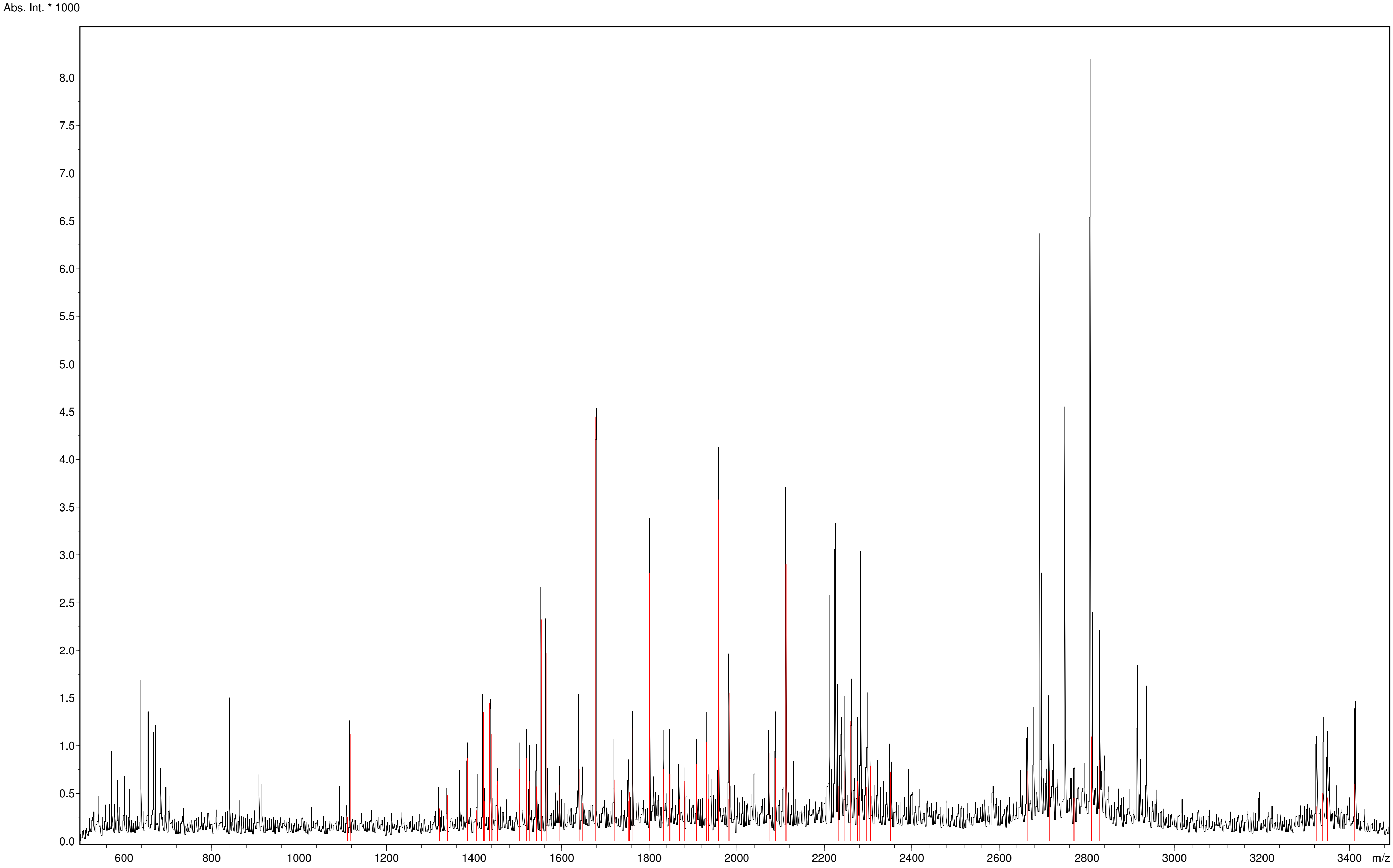



Copyright © 1999-2008 Bruker Daltonik GmbH

#### Spectrum Analysis Report
### Cytochrome c1

**Sequence data:**

Intensity Coverage: 2.1 % (0 cnts)
Sequence Coverage MS: 0.0%
Sequence Coverage MS/MS: 0.0%
pI (isoelectric point): 9.0

| | | | | | | | | | | | |
|---|---|---|---|---|---|---|---|---|---|---|---|
| 10 | 20 | 30 | 40 | 50 | 60 | 70 | 80 | 90 | 100 | 110 | 120 |
| MFSNLSKRWA | QRTLSKSFYS | TATGAASKSG | KLTQKLVTAG | VAAAGITAST | LLYADSLTAE | AMTAAEHGLH | APAYAWSHNG | PFETFDHASI | RRGYQVYREV | CAACHSLDRV | AWRTLVGVSH |
| 130 | 140 | 150 | 160 | 170 | 180 | 190 | 200 | 210 | 220 | 230 | 240 |
| TNEEVRNMAE | EFEYDDEPDE | QGNPKKRPGK | LSDYIPGPYP | NEQAARAANQ | GALPPDLSLI | VKARHGGCDY | IFSLLTGYPD | EPPAGVALPP | GSNYNPYFPG | GSIAMARVLF | DDMVEYEDGT |
| 250 | 260 | 270 | 280 | 290 | 300 | 310 | | | | | |
| PATTSQMAKD | VTTFLNWCAE | PEHDERKRLG | LKTVIILSSL | YLLSIWVKKF | KWAGIKTRKF | VFNPPKPRK | | | | | |

**Display Parameter:**

MH+ (mono): 1.008 MH+ (avg): 1.008 Threshold (a.i.): 0.000 Tolerance (Da): 0.500
Number of Peaks: 59

**Peaklist:**

| Peak | Mass | Intensity | Peak | Mass | Intensity | Peak | Mass | Intensity | Peak | Mass | Intensity | Peak | Mass | Intensity |
|---|---|---|---|---|---|---|---|---|---|---|---|---|---|---|
| 1 | 1110.054 | 251.933 | 2 | 1116.666 | 1120.728 | 3 | 1320.786 | 336.923 | 4 | 1338.796 | 478.042 | 5 | 1367.800 | 492.717 |
| 6 | 1384.757 | 867.658 | 7 | 1405.859 | 322.220 | 8 | 1420.736 | 1354.999 | 9 | 1423.900 | 417.522 | 10 | 1435.803 | 1446.506 |
| 11 | 1438.888 | 1118.738 | 12 | 1442.879 | 395.058 | 13 | 1453.847 | 631.210 | 14 | 1502.896 | 753.977 | 15 | 1519.813 | 867.060 |
| 16 | 1525.886 | 624.680 | 17 | 1541.791 | 570.619 | 18 | 1553.780 | 2322.852 | 19 | 1563.931 | 1966.513 | 20 | 1595.910 | 586.078 |
| 21 | 1639.888 | 753.983 | 22 | 1646.910 | 397.868 | 23 | 1677.978 | 4447.644 | 24 | 1720.041 | 642.049 | 25 | 1751.994 | 415.628 |
| 26 | 1754.932 | 510.120 | 27 | 1763.052 | 1182.452 | 28 | 1800.975 | 2805.140 | 29 | 1832.154 | 756.133 | 30 | 1847.101 | 706.413 |
| 31 | 1869.060 | 431.945 | 32 | 1880.092 | 627.140 | 33 | 1908.071 | 807.081 | 34 | 1930.053 | 1035.089 | 35 | 1935.077 | 437.616 |
| 36 | 1958.155 | 3577.306 | 37 | 1980.139 | 437.091 | 38 | 1984.125 | 1555.893 | 39 | 2073.124 | 924.758 | 40 | 2089.122 | 866.665 |
| 41 | 2112.212 | 2898.941 | 42 | 2233.129 | 577.860 | 43 | 2247.157 | 736.130 | 44 | 2260.204 | 1257.258 | 45 | 2276.220 | 623.620 |
| 46 | 2279.220 | 471.446 | 47 | 2296.234 | 563.204 | 48 | 2305.202 | 783.002 | 49 | 2351.187 | 719.146 | 50 | 2663.405 | 733.583 |
| 51 | 2713.317 | 733.201 | 52 | 2770.336 | 451.211 | 53 | 2810.355 | 1092.892 | 54 | 2829.358 | 848.095 | 55 | 2936.554 | 661.985 |
| 56 | 3323.786 | 360.067 | 57 | 3338.798 | 505.468 | 58 | 3348.667 | 454.129 | 59 | 3411.760 | 600.095 | | | |


Copyright © 1999-2008 Bruker Daltonik GmbH

**Spectrum Analysis Report**

#### Rieske subunit

| | | | | | | | |
|---|---|---|---|---|---|---|---|
| Sequence Name: | | Formula: | | Parentmass: | | Mass Error: | |
| MH+ (mono): | 1.008 | MH+ (avg): | 1.008 | Threshold (a.i.): | 0.000 | Tolerance (Da): | 0.500 |
| Number of Peaks: | 59 | Above Threshold: | | Assigned Peaks: | | Not assigned Peaks: | |

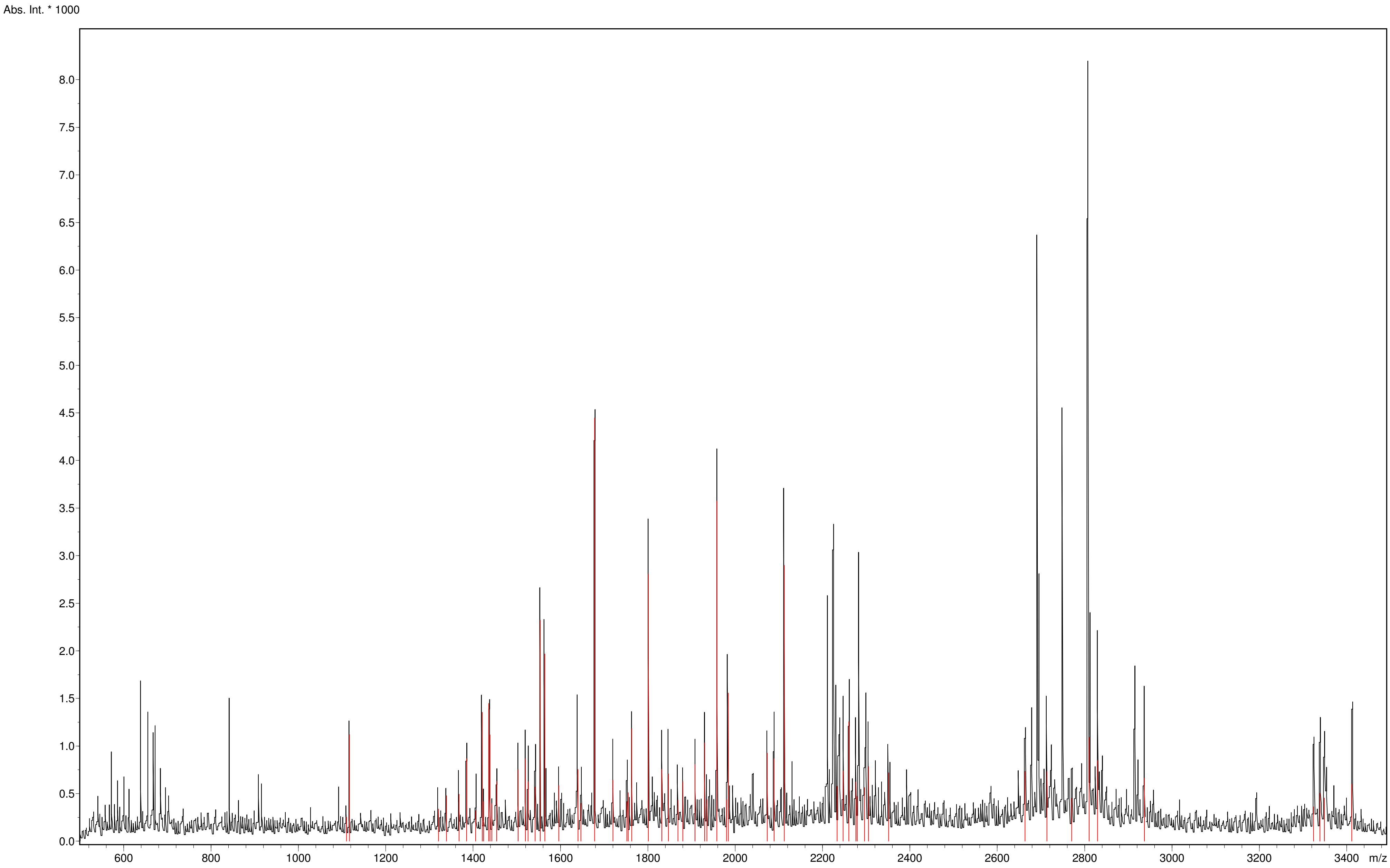



Copyright © 1999-2008 Bruker Daltonik GmbH

#### Spectrum Analysis Report

### Rieske subunit

**Sequence data:**

| | | | |
|---|---|---|---|
| Intensity Coverage: | 2.1 % (1182 cnts) | Sequence Coverage MS: | 7.9% |
| Sequence Coverage MS/MS: | 0.0% | pI (isoelectric point): | 9.3 |

| 10 | 20 | 30 | 40 | 50 | 60 | 70 | 80 | 90 | 100 | 110 | 120 |
|---|---|---|---|---|---|---|---|---|---|---|---|
| MLGIRSSVKT | CFKPMSLTSK | RLISQSLLAS | KSTYRTPNFD | DVLKENNDAD | KGRSYAYFMV | GAMGLLSSAG | AKSTVETFIS | SMTATADVLA | MAKVEVNLAA | IPLGKNVVVK | WQGKPVFIRH |
| 130 | 140 | 150 | 160 | 170 | 180 | 190 | 200 | 210 | 220 | | |
| RTPHEIQEAN | SVDMSALKDP | QTDADRVKDP | QWLIMLGICT | HLGCVPIGEA | GDFGGWFCPC | HGSHYDISGR | IRKGPAPLNL | EIPAYEFDGD | KVIVG | | |

**Display Parameter:**

| | | | | | | | |
|---|---|---|---|---|---|---|---|
| MH+ (mono): | 1.008 | MH+ (avg): | 1.008 | Threshold (a.i.): | 0.000 | Tolerance (Da): | 0.500 |
| Number of Peaks: | 59 | | | | | | |

**Peaklist:**

| Peak | Mass | Intensity | Peak | Mass | Intensity | Peak | Mass | Intensity | Peak | Mass | Intensity | Peak | Mass | Intensity |
|---|---|---|---|---|---|---|---|---|---|---|---|---|---|---|
| 1 | 1110.054 | 251.933 | 2 | 1116.666 | 1120.728 | 3 | 1320.786 | 336.923 | 4 | 1338.796 | 478.042 | 5 | 1367.800 | 492.717 |
| 6 | 1384.757 | 867.658 | 7 | 1405.859 | 322.220 | 8 | 1420.736 | 1354.999 | 9 | 1423.900 | 417.522 | 10 | 1435.803 | 1446.506 |
| 11 | 1438.888 | 1118.738 | 12 | 1442.879 | 395.058 | 13 | 1453.847 | 631.210 | 14 | 1502.896 | 753.977 | 15 | 1519.813 | 867.060 |
| 16 | 1525.886 | 624.680 | 17 | 1541.791 | 570.619 | 18 | 1553.780 | 2322.852 | 19 | 1563.931 | 1966.513 | 20 | 1595.910 | 586.078 |
| 21 | 1639.888 | 753.983 | 22 | 1646.910 | 397.868 | 23 | 1677.978 | 4447.644 | 24 | 1720.041 | 642.049 | 25 | 1751.994 | 415.628 |
| 26 | 1754.932 | 510.120 | 27 | 1763.052 | 1182.452 | 28 | 1800.975 | 2805.140 | 29 | 1832.154 | 756.133 | 30 | 1847.101 | 706.413 |
| 31 | 1869.060 | 431.945 | 32 | 1880.092 | 627.140 | 33 | 1908.071 | 807.081 | 34 | 1930.053 | 1035.089 | 35 | 1935.077 | 437.616 |
| 36 | 1958.155 | 3577.306 | 37 | 1980.139 | 437.091 | 38 | 1984.125 | 1555.893 | 39 | 2073.124 | 924.758 | 40 | 2089.122 | 866.665 |
| 41 | 2112.212 | 2898.941 | 42 | 2233.129 | 577.860 | 43 | 2247.157 | 736.130 | 44 | 2260.204 | 1257.258 | 45 | 2276.220 | 623.620 |
| 46 | 2279.220 | 471.446 | 47 | 2296.234 | 563.204 | 48 | 2305.202 | 783.002 | 49 | 2351.187 | 719.146 | 50 | 2663.405 | 733.583 |
| 51 | 2713.317 | 733.201 | 52 | 2770.336 | 451.211 | 53 | 2810.355 | 1092.892 | 54 | 2829.358 | 848.095 | 55 | 2936.554 | 661.985 |
| 56 | 3323.786 | 360.067 | 57 | 3338.798 | 505.468 | 58 | 3348.667 | 454.129 | 59 | 3411.760 | 600.095 | | | |


Copyright © 1999-2008 Bruker Daltonik GmbH

**Spectrum Analysis Report**

#### iso-1 Cc

| | | | | | | | |
|---|---|---|---|---|---|---|---|
| Sequence Name: | | Formula: | | Parentmass: | | Mass Error: | |
| MH+ (mono): | 1.008 | MH+ (avg): | 1.008 | Threshold (a.i.): | 0.000 | Tolerance (Da): | 0.500 |
| Number of Peaks: | 59 | Above Threshold: | | Assigned Peaks: | | Not assigned Peaks: | |

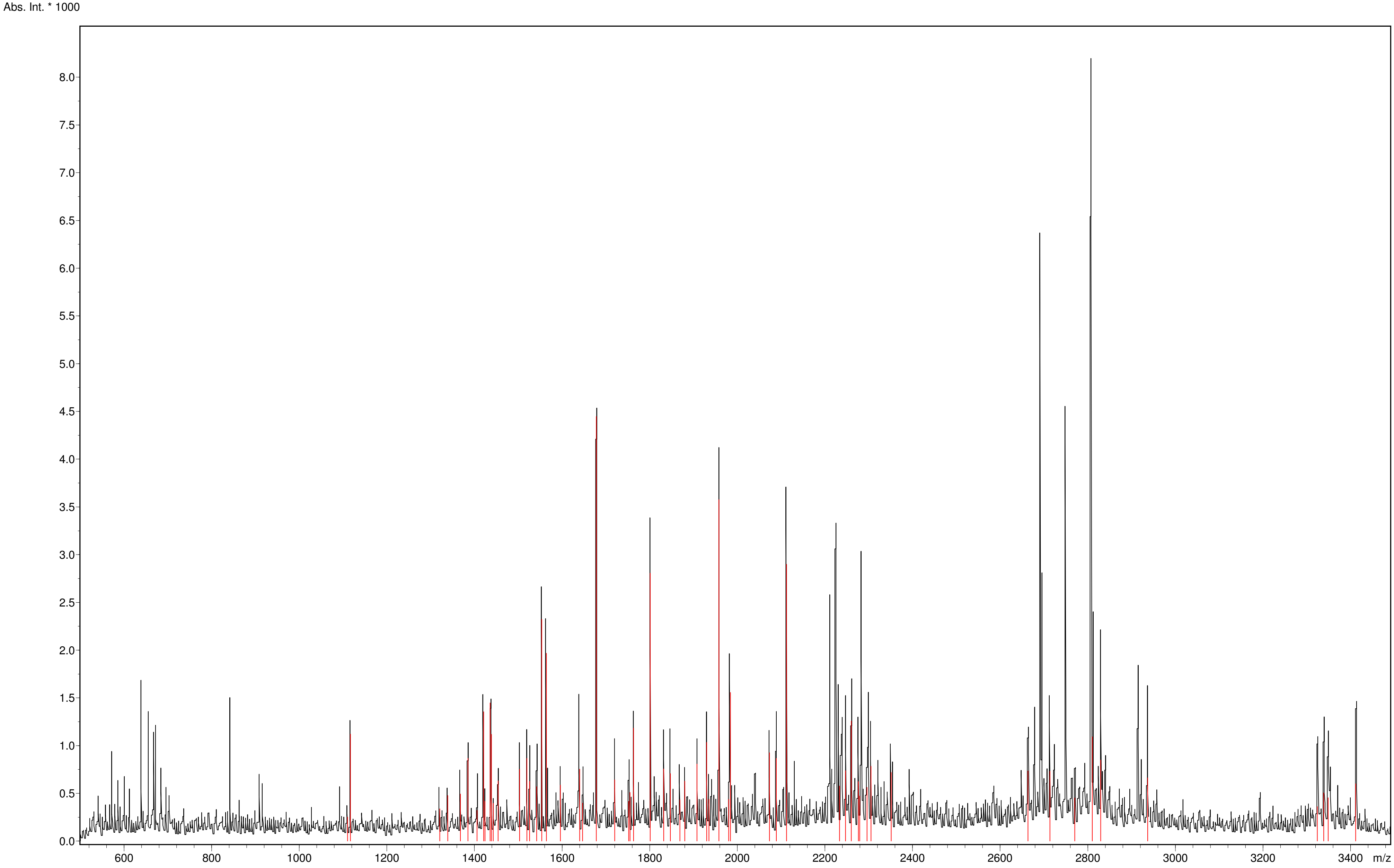




Copyright © 1999-2008 Bruker Daltonik GmbH

#### Spectrum Analysis Report

### iso-1 Cc

**Sequence data:**

| | | | |
|---|---|---|---|
| Intensity Coverage: | 2.9 % (0 cnts) | Sequence Coverage MS: | 0.0% |
| Sequence Coverage MS/MS: | 0.0% | pI (isoelectric point): | 10.1 |

| 10 | 20 | 30 | 40 | 50 | 60 | 70 | 80 | 90 | 100 | 110 |
|---|---|---|---|---|---|---|---|---|---|---|
| MTEFKAGSAK | KGATLFKTRC | LQCHTVEKGG | PHKVGPNLHG | IFGRHSGQAE | GYSYTDANIK | KNVLWDENNM | SEYLTNPKKY | IPGTKMAFGG | LKKEKDRNDL | ITYLKKACE |

**Display Parameter:**

| | | | | | | | |
|---|---|---|---|---|---|---|---|
| MH+ (mono): | 1.008 | MH+ (avg): | 1.008 | Threshold (a.i.): | 0.000 | Tolerance (Da): | 0.500 |
| Number of Peaks: | 59 | | | | | | |

**Peaklist:**

| Peak | Mass | Intensity | Peak | Mass | Intensity | Peak | Mass | Intensity | Peak | Mass | Intensity | Peak | Mass | Intensity |
|---|---|---|---|---|---|---|---|---|---|---|---|---|---|---|
| 1 | 1110.054 | 251.933 | 2 | 1116.666 | 1120.728 | 3 | 1320.786 | 336.923 | 4 | 1338.796 | 478.042 | 5 | 1367.800 | 492.717 |
| 6 | 1384.757 | 867.658 | 7 | 1405.859 | 322.220 | 8 | 1420.736 | 1354.999 | 9 | 1423.900 | 417.522 | 10 | 1435.803 | 1446.506 |
| 11 | 1438.888 | 1118.738 | 12 | 1442.879 | 395.058 | 13 | 1453.847 | 631.210 | 14 | 1502.896 | 753.977 | 15 | 1519.813 | 867.060 |
| 16 | 1525.886 | 624.680 | 17 | 1541.791 | 570.619 | 18 | 1553.780 | 2322.852 | 19 | 1563.931 | 1966.513 | 20 | 1595.910 | 586.078 |
| 21 | 1639.888 | 753.983 | 22 | 1646.910 | 397.868 | 23 | 1677.978 | 4447.644 | 24 | 1720.041 | 642.049 | 25 | 1751.994 | 415.628 |
| 26 | 1754.932 | 510.120 | 27 | 1763.052 | 1182.452 | 28 | 1800.975 | 2805.140 | 29 | 1832.154 | 756.133 | 30 | 1847.101 | 706.413 |
| 31 | 1869.060 | 431.945 | 32 | 1880.092 | 627.140 | 33 | 1908.071 | 807.081 | 34 | 1930.053 | 1035.089 | 35 | 1935.077 | 437.616 |
| 36 | 1958.155 | 3577.306 | 37 | 1980.139 | 437.091 | 38 | 1984.125 | 1555.893 | 39 | 2073.124 | 924.758 | 40 | 2089.122 | 866.665 |
| 41 | 2112.212 | 2898.941 | 42 | 2233.129 | 577.860 | 43 | 2247.157 | 736.130 | 44 | 2260.204 | 1257.258 | 45 | 2276.220 | 623.620 |
| 46 | 2279.220 | 471.446 | 47 | 2296.234 | 563.204 | 48 | 2305.202 | 783.002 | 49 | 2351.187 | 719.146 | 50 | 2663.405 | 733.583 |
| 51 | 2713.317 | 733.201 | 52 | 2770.336 | 451.211 | 53 | 2810.355 | 1092.892 | 54 | 2829.358 | 848.095 | 55 | 2936.554 | 661.985 |
| 56 | 3323.786 | 360.067 | 57 | 3338.798 | 505.468 | 58 | 3348.667 | 454.129 | 59 | 3411.760 | 600.095 | | | |


Copyright © 1999-2008 Bruker Daltonik GmbH

#### Spectrum Analysis Report

### iso-2 Cc

| | | | | | | | |
|---|---|---|---|---|---|---|---|
| Sequence Name: | | Formula: | | Parentmass: | | Mass Error: | |
| MH+ (mono): | 1.008 | MH+ (avg): | 1.008 | Threshold (a.i.): | 0.000 | Tolerance (Da): | 0.500 |
| Number of Peaks: | 59 | Above Threshold: | | Assigned Peaks: | | Not assigned Peaks: | |

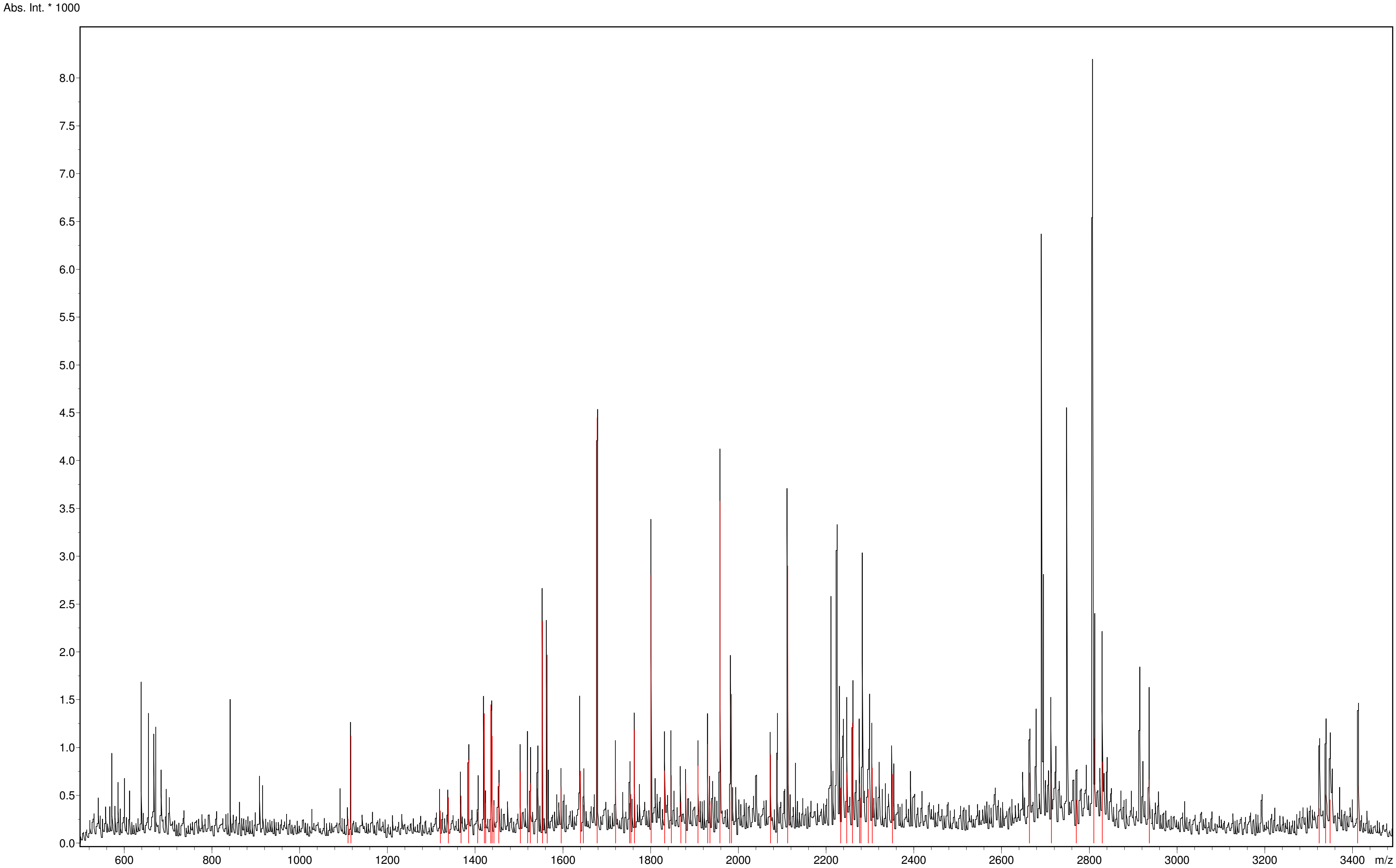



Copyright © 1999-2008 Bruker Daltonik GmbH

### Spectrum Analysis Report
#### iso-2 Cc

**Sequence data:**

| | | | |
|---|---|---|---|
| Intensity Coverage: | 2.9 % (1622 cnts) | Sequence Coverage MS: | 12.4% |
| Sequence Coverage MS/MS: | 0.0% | pI (isoelectric point): | 10.2 |

```
    10         20         30         40         50         60         70         80         90         100        110        120
MAKESTGFKP GSAKKGATLF KTRCQQCHTI EEGGPNKVGP NLHGIFGRHS GQVKGYSYTD ANINKNVKWD EDSMSEYLTN PKKYIPGTKM AFAGLKKEKD RNDLITYMTK AAK
```

**Display Parameter:**

| | | | | | | | |
|---|---|---|---|---|---|---|---|
| MH+ (mono): | 1.008 | MH+ (avg): | 1.008 | Threshold (a.i.): | 0.000 | Tolerance (Da): | 0.500 |
| Number of Peaks: | 59 | | | | | | |

**Peaklist:**

| Peak | Mass | Intensity | Peak | Mass | Intensity | Peak | Mass | Intensity | Peak | Mass | Intensity | Peak | Mass | Intensity |
|---|---|---|---|---|---|---|---|---|---|---|---|---|---|---|
| 1 | 1110.054 | 251.933 | 2 | 1116.666 | 1120.728 | 3 | 1320.786 | 336.923 | 4 | 1338.796 | 478.042 | 5 | 1367.800 | 492.717 |
| 6 | 1384.757 | 867.658 | 7 | 1405.859 | 322.220 | 8 | 1420.736 | 1354.999 | 9 | 1423.900 | 417.522 | 10 | 1435.803 | 1446.506 |
| 11 | 1438.888 | 1118.738 | 12 | 1442.879 | 395.058 | 13 | 1453.847 | 631.210 | 14 | 1502.896 | 753.977 | 15 | 1519.813 | 867.060 |
| 16 | 1525.886 | 624.680 | 17 | 1541.791 | 570.619 | 18 | 1553.780 | 2322.852 | 19 | 1563.931 | 1966.513 | 20 | 1595.910 | 586.078 |
| 21 | 1639.888 | 753.983 | 22 | 1646.910 | 397.868 | 23 | 1677.978 | 4447.644 | 24 | 1720.041 | 642.049 | 25 | 1751.994 | 415.628 |
| 26 | 1754.932 | 510.120 | 27 | 1763.052 | 1182.452 | 28 | 1800.975 | 2805.140 | 29 | 1832.154 | 756.133 | 30 | 1847.101 | 706.413 |
| 31 | 1869.060 | 431.945 | 32 | 1880.092 | 627.140 | 33 | 1908.071 | 807.081 | 34 | 1930.053 | 1035.089 | 35 | 1935.077 | 437.616 |
| 36 | 1958.155 | 3577.306 | 37 | 1980.139 | 437.091 | 38 | 1984.125 | 1555.893 | 39 | 2073.124 | 924.758 | 40 | 2089.122 | 866.665 |
| 41 | 2112.212 | 2898.941 | 42 | 2233.129 | 577.860 | 43 | 2247.157 | 736.130 | 44 | 2260.204 | 1257.258 | 45 | 2276.220 | 623.620 |
| 46 | 2279.220 | 471.446 | 47 | 2296.234 | 563.204 | 48 | 2305.202 | 783.002 | 49 | 2351.187 | 719.146 | 50 | 2663.405 | 733.583 |
| 51 | 2713.317 | 733.201 | 52 | 2770.336 | 451.211 | 53 | 2810.355 | 1092.892 | 54 | 2829.358 | 848.095 | 55 | 2936.554 | 661.985 |
| 56 | 3323.786 | 360.067 | 57 | 3338.798 | 505.468 | 58 | 3348.667 | 454.129 | 59 | 3411.760 | 600.095 | | | |


Copyright © 1999-2008 Bruker Daltonik GmbH
