## Supplementary Figures and Tables for "Unveiling the role of yeast cytochrome *c* isoforms in the assembly of mitochondrial supercomplexes and the control of respiratory chain rate"

**APPENDIX A. SUPPLEMENTARY FIGURES AND TABLES**

Alejandra Guerra-Castellano[a,*], Manuel Aneas[a], Joaquín Tamargo-Azpilicueta[a], Inmaculada Márquez[b], José Luis Olloqui-Sariego[b], Juan José Calvente[b], Miguel A. De la Rosa[a], Irene Díaz-Moreno[a,*]

[a]Instituto de Investigaciones Químicas, Centro de Investigaciones Científicas Isla de la Cartuja, Universidad de Sevilla - CSIC. Avda. Americo Vespucio 49, 41092 Sevilla, Spain.
[b]Departamento de Química Física, Universidad de Sevilla, Profesor García González 1, 41012 Sevilla, Spain.

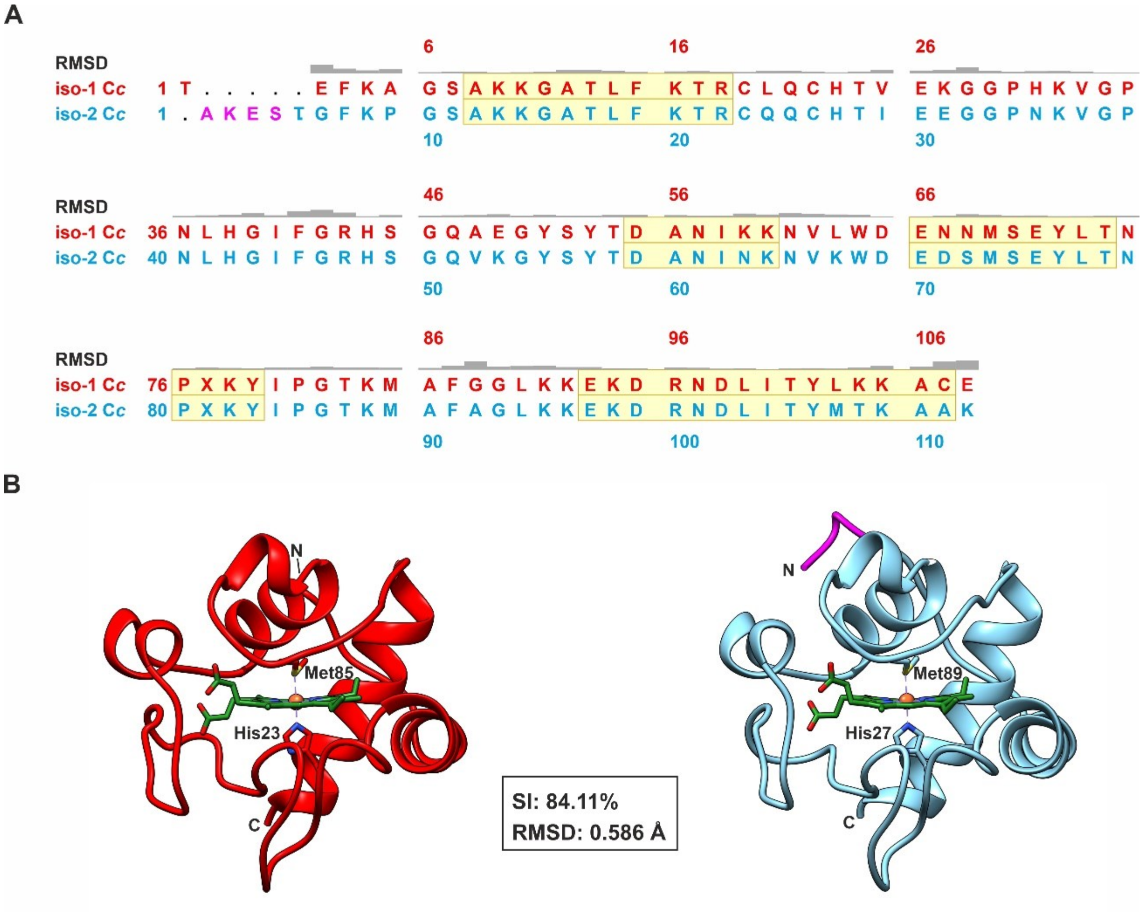


**Figure S1. Sequence alignment and 3D structural models of yeast cytochrome *c* isoforms**. (**A**) Alignment between iso-1 (red) and iso-2 (blue) sequences of yeast C*c*. RMSD of the backbone is indicated per residue. The sequence highlighted in yellow indicates α-helix structure regions. X residue: Lysine Trimethyllysine. (**B**) Ribbon representation of tridimensional structures of yeast C*c* isoforms. The iso-1 C*c* (PDB 3CX5; Solmaz & Hunte, 2008) and iso-2 C*c* (PDB 3CXH; Solmaz & Hunte, 2008) are represented in red and blue, respectively. The four amino acid extension present at the N-terminus of iso-2 C*c* is highlighted in magenta. The heme group is colored green and the iron atom is orange. The axial ligands and the N and C-terminus are indicated. SI. Sequence identity; RMSD: Root-Mean-Square Deviation. The figure has been generated with the software Chimera (Pettersen *et al*., 2004).

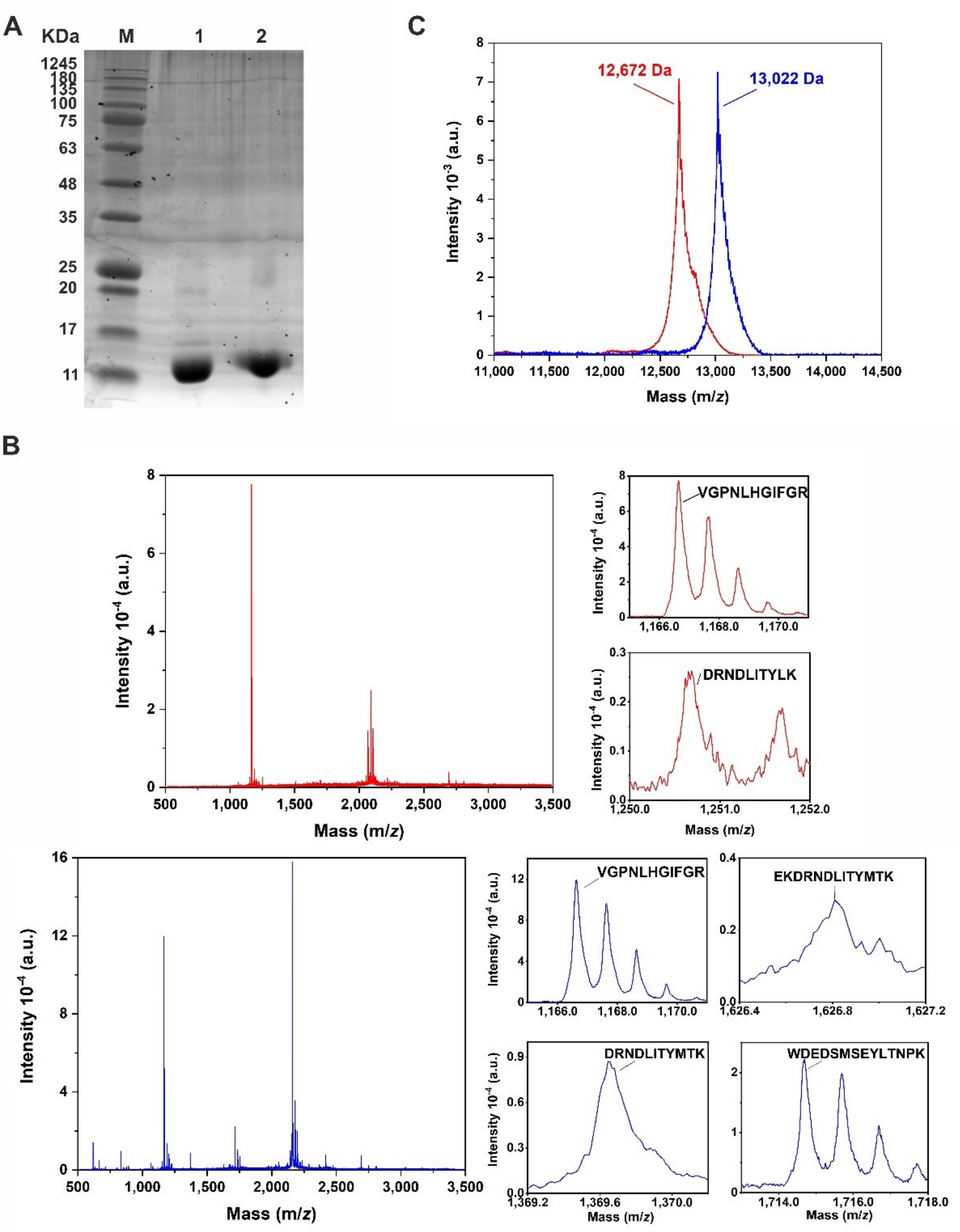


**Figure S2. Yeast cytochrome *c* isoforms characterization.** (**A**) SDS-PAGE of purified C*c* recombinant iso-1 (lane 1) and iso-2 (lane 2). M: molecular weight marker. (**B**) Tryptic digestion analysis of C*c* isoforms 1 (top, red) and 2 (bottom, blue). The right panels display the different detected peptides in the mass spectra and their corresponding sequences. (**C**) MALDI-TOF mass spectra of C*c* isoforms 1 (red) and 2 (blue). Isoforms masses are indicated in the same color pattern.

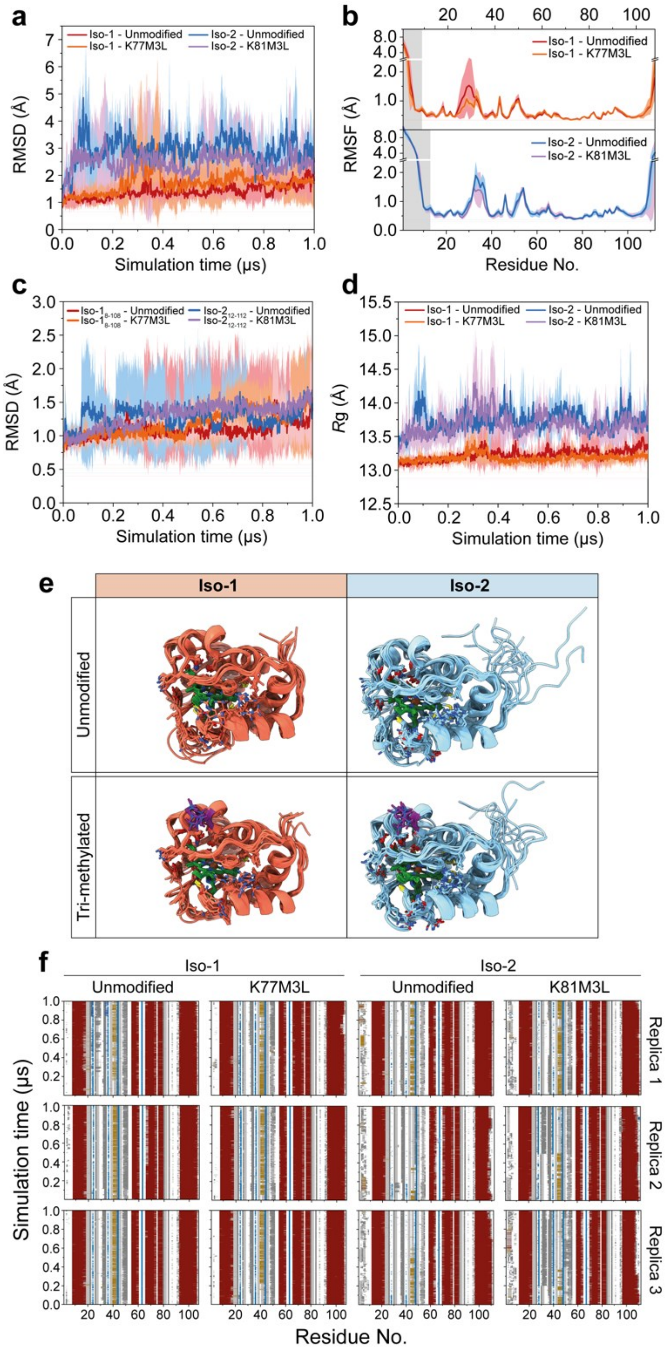


**Figure S3. Molecular dynamics simulation of iso-1 and iso-2 cytochrome *c*. (A)** Backbone root-mean-square deviation (RMSD) analysis of the C*c* isoforms 1 μs trajectories using the initial structure as the reference. The standard deviation for n=3 replicas is represented as shadowed lines. **(B)** Per-residue backbone atom root-mean square fluctuations (RMSF) during the trajectories of iso-1 (upper panel) and iso-2 (lower pannel) C*c*. (C) RMSD analysis of the folded region of C*c* iso-1 and iso-2 during 1 μs trajectories. Shadowed areas in grey in panel **(B)** indicate the flexible areas excluded for this analysis (i.e., residues 1-8 in iso-1 and 1-12 in iso-2). **(D)** Evolution of the radius of gyration of the full length protein along the trajectories. **(E)** 100 ns protein snapshots of the first replica of each construct analyzed. The heme group was depicted in green and tri-methyl lysine in purple. **(F)** Secondary structure analysis of each trajectory, according to DSSP method (Kabsch & Sander, 1983). $\alpha$-helices were depicted in red; $3_{10}$ helices in yellow, $\beta$-sheets in blue, turns in grey and coil regions in white.

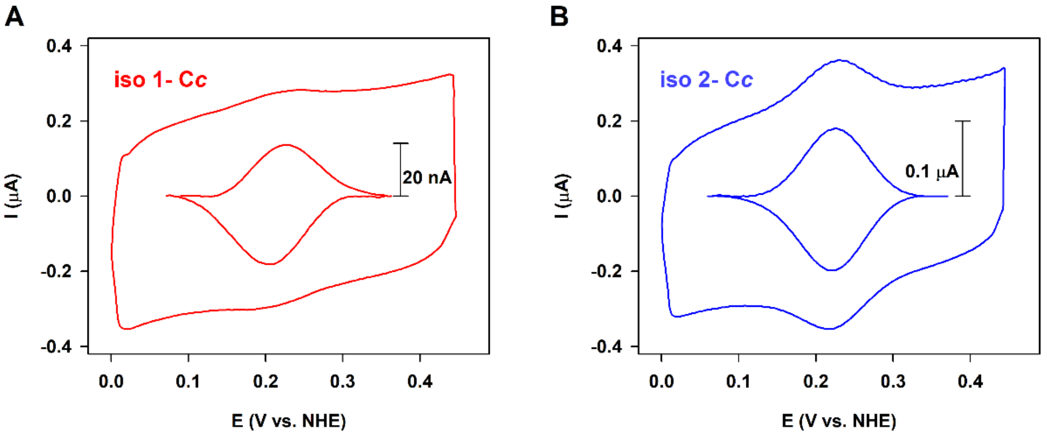


**Figure S4. Voltammetric characterization of the interfacial electron transfer of yeast cytochrome *c* isoforms. (A)** Cyclic voltammograms of iso-1 and **(B)** iso-2 C*c* immobilized onto a mixed-SAM-modified gold electrode and measured in 20 mM sodium phosphate buffer at pH 7.0 at 25 °C.

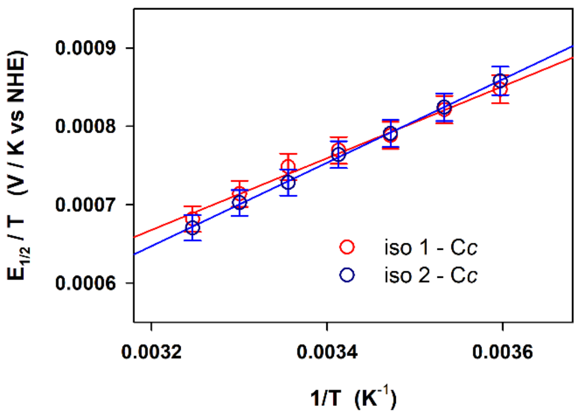


**Figure S5. Electrochemical characterization of yeast cytochrome *c* isoforms with temperature.** Variation of the temperature normalized midpoint potential $E_{1/2}/T$, and with the temperature for isoforms 1 (red symbols) and 2 (blue symbols) immobilized onto a mixed-SAM-modified gold electrode and measured in 20 mM sodium phosphate buffer at pH 7.0. Solid lines correspond to the linear least-square fits of experimental data.

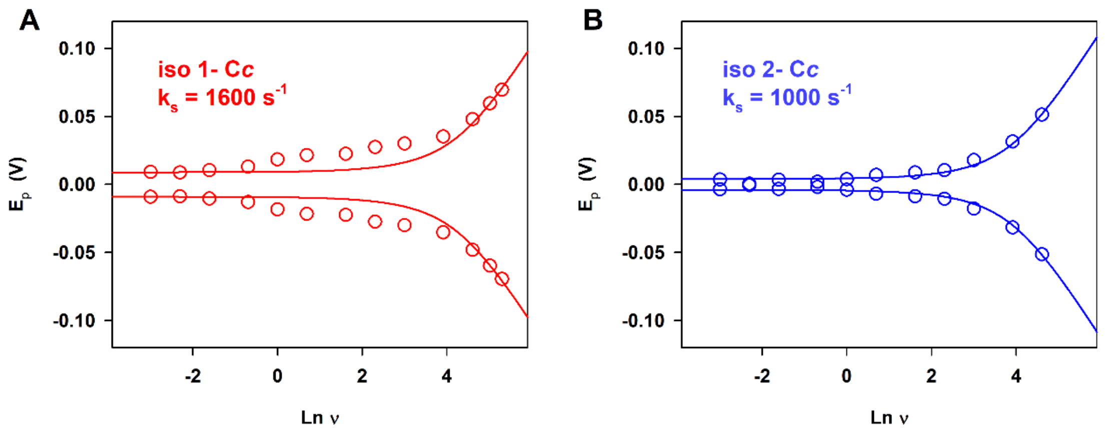


**Figure S6. Kinetic characterization of the interfacial electron transfer of yeast cytochrome *c* isoforms. (A)** Trumpet plots obtained from the variation of anodic and cathodic peak potentials with the logarithm of the potential scan rate of iso-1 and **(B)** iso-2 C*c* immobilized onto a mixed-SAM-modified gold electrode and measured in 20 mM sodium phosphate buffer at pH 7.0 at 25 ºC.

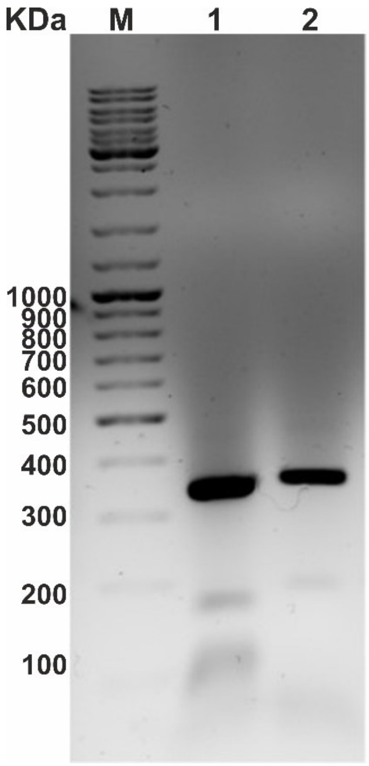


**Figure S7. Identification of *cyc1* and *cyc7* in *Saccharomyces cerevisiae* complementary strains.** Lanes correspond to the amplificated DNA from *cyc1*$^{+}$ (lane 1) and *cyc7*$^{+}$ (lane 2) strains. The obtained bands correspond to *cyc1* and *cyc7* based on their molecular weights (330 bp and 342 bp, respectively). M: molecular weight marker; *cyc1*$^{+}$: iso-1 C*c* complementary strain; *cyc7*$^{+}$: iso-2 C*c* complementary strain.

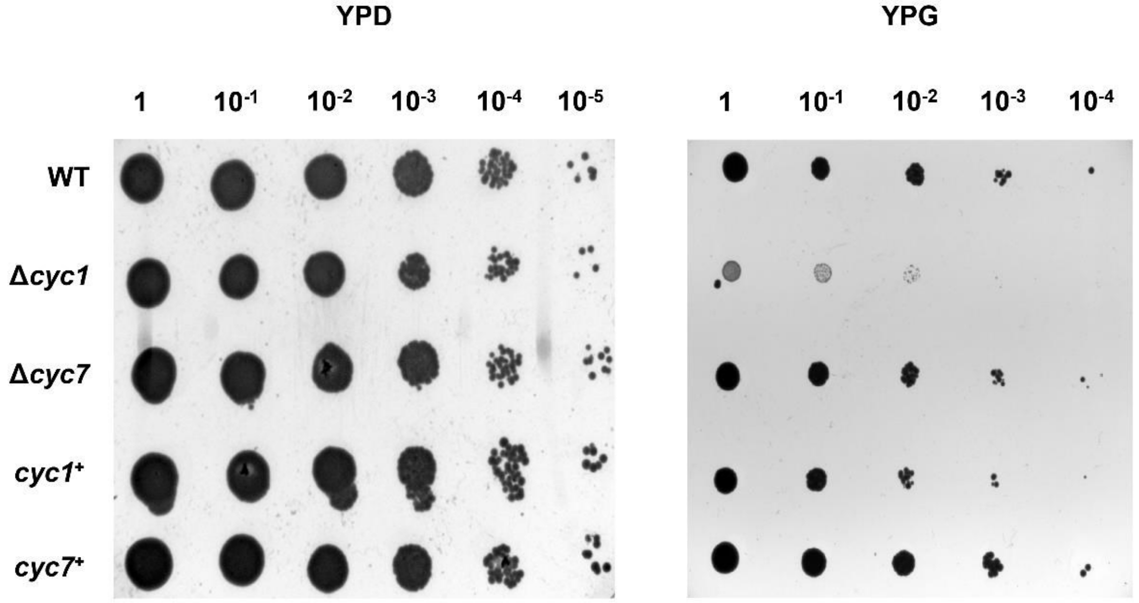


**Figure S8. Growth patterns of *Saccharomyces cerevisiae* strains.** Yeast strains were grown in YPD and YPG media from an OD = 0.5 (equivalent to $0.75{\cdot}10^7$ cells; Feldmann, 2010) and making dilutions described in the horizontal axis. Both media plates were incubated for 24 h at 30 °C. WT: wild-type, Δ*cyc1*: iso-1 C*c* deficient strain, Δ*cyc7*: iso-2 C*c* deficient strain, *cyc1*$^+$: iso-1 C*c* complementary strain, and *cyc7*$^+$: iso-2 C*c* complementary strain.

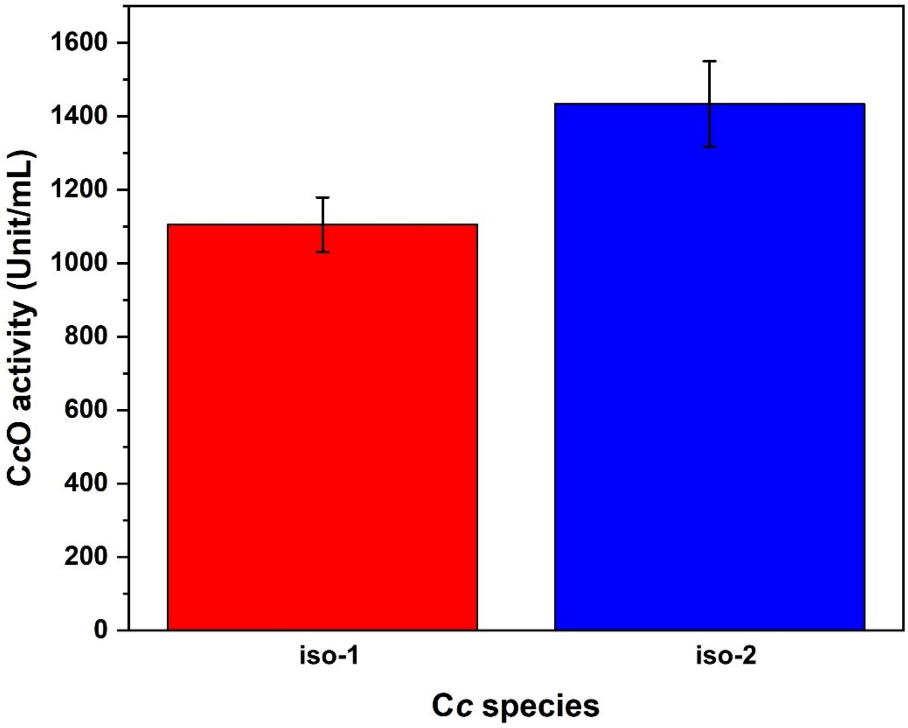


**Figure S9. Cytochrome *c* oxidase activity.** Cytochrome *c* oxidase (C*c*O) activity of isolated CIV upon addition of exogenous iso-1 (red bars) or iso-2 C*c* (blue bars). All data represent at least the mean ±SD of three independent experiments.

**Table S1. Primers for extraction of *cyc1* and *cyc7***

| | |
|---|---|
| To extract *cyc1* from the genomic DNA of *S. cerevisiae* | |
| pBTR1-*cyc1-fw* | 5'-GAAGGAGATATATCCATGACTGAATTCAAGGCCGGT-3' |
| pBTR1-*cyc1-rv* | 5'-AGCTTGGCCGGATCCTTACTCACAGGCTTTTTTCAA-3' |
| To extract *cyc7* from the genomic DNA of *S. cerevisiae* | |
| pBTR1-*cyc7-fw* | 5'-GAAGGAGATATATCCATGGCTAAAGAAAGTACGGGATTCAAAC-3' |
| pBTR1-*cyc7-rv* | 5'-AGCTTGGCCGGATCCCTATTTGGCAGCCTTTGTCAT-3' |

**Table S2. Primers utilized to obtain complementary strains**

| | |
|---|---|
| To linearize pCM189 plasmid | |
| *fw* Pst1 | 3'-ctgcaggagggccgcatcatgtaattag-5' |
| *rv* BamHI | 3'-ggatcccccgaattgatccggtaatttag-5' |
| To obtain the inserts of the *cyc1* and *cyc7* from pBTR1 plasmid | |
| *fw-cyc1* | 3'-caattcgggggatccATGACTGAATTCAAGGCCGGT-5' |
| *rv-cyc1* | 3'-gcggccctcctgcagTTACTCACAGGCTTTTTTCAA-5' |
| *fw-cyc7* | 3'-caattcgggggatccATGGCTAAAGAAAGTACGGGA-5' |
| *rv-cyc7* | 3'-gcggccctcctgcagCTATTTGGCAGCCTTTGTCATATAAGT-5' |

**Table S3. Distances for the transfer between CIII-C*c*-CIV**

| Interatomic distance | C*c* isoform | $III_2IV$ (Å) | $III_2IV_2$ (Å) |
|---|---|---|---|
| Fe-heme C$c_1$ / $Cu_A$-site CIV distance | - | 62.938 | 61.896 |
| Fe-heme C*c* / Fe-heme C$c_1$ distance | iso-1 | 17.687 | |
| | iso-2 | 17.591 | 17.338 |
| Fe-heme C*c* (CIII)/ Fe-heme C*c* distance (CIV) | iso-1 | 69.325 | |
| | iso-2 | 69.240 | 69.240 |
| Fe-heme $C_c$ / $Cu_A$-site CIV distance | iso-1 | 21.463 | |
| | iso-2 | 21.357 | 22.730 |

C*c*: cytochrome *c*; C$c_1$: cytochrome $c_1$; CIII: complex III; CIV: complex IV; $III_2IV$: supercomplex consists of a CIII dimer with a CIV; $III_2IV_2$: supercomplex consists of a CIII dimer flanked by two CIV.
